# A Thermodynamically Consistent Reaction–Diffusion Model of Spatial Proofreading in One and Two Dimensions

**DOI:** 10.64898/2026.07.28.741167

**Authors:** Tommaso Rossi, Thibault Fillion, Francesco Piazza

## Abstract

Spatial proofreading is a mechanism that can enhance molecular discrimination by exploiting nonequilibrium diffusive transport between spatially separated source and readout regions. Here, we introduce a thermodynamically consistent reaction–diffusion model in which a gradient of active substrates is generated self-consistently by a reversible kinase–phosphatase switch coupled to nucleotide chemostats. The resulting chemical-potential gradient explicitly controls the nonequilibrium driving and allows the discrimination potential to be related to microscopic chemical rates and diffusive transport. A simple one-dimensional analysis shows that the ability of an enzyme to discriminate between wrong and right substrates is governed by a subtle balance between substrate-gradient confinement, controlled by phosphatase activity, the diffusive crossing time across the source–readout domain, and selective complex dissociation. We then extend the model to two-dimensional domains, showing that source, i.e. kinase, localization modulates spatial specificity by shaping the effective diffusive paths to the readout boundary. Finally, stochastic simulations reveal that molecular fluctuations generate intermittent wrong-readout events, which can be characterized through an event-weighted measure of specificity. Interestingly, we find that fluctuations promote frequent transitions to long-residence states in which discrimination is better than predicted by the deterministic estimate. Overall, our work highlights the importance of thermodynamically consistent descriptions of spatial proofreading and clarifies how energy input, transport, and spatial organization jointly shape biochemical discrimination.

## 1. Introduction

The ability to discriminate between correct and wrong substrates is fundamental in many biological processes based on the formation of molecular complexes, from translation to signaling networks. In this work, we consider a proofreading enzyme (E) that can bind to two alternative substrates, a correct (R) and a wrong (W) one, present in the system at much higher concentration. These can form the complexes *ER* and *EW*, respectively. Discrimination is quantified through the specificity

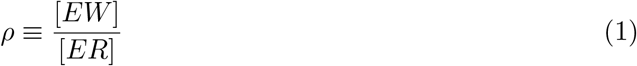

where smaller values of *ρ* correspond to higher accuracy. At thermodynamic equilibrium, specificity is constrained by the free-energy difference between the correct and wrong complexes,

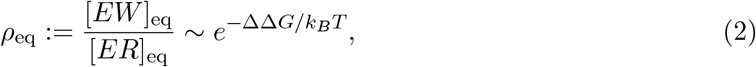

with ΔΔ*G* = Δ*G*_W_ *−* Δ*G*_R_. However, in many biological systems this difference is of the order of only a few *k*_*B*_*T*. The equilibrium limit associated with this energetic scale is therefore not sufficient to explain the high levels of accuracy observed experimentally: in translation, error rates are of the order of ~10^*−*4^ per codon [1], whereas in DNA replication the final error probability can decrease down to ~10^*−*10^ per base [2]. These orders of magnitude indicate the presence of nonequilibrium discrimination mechanisms that allow biological systems to overcome the limit imposed by thermodynamic equilibrium. These processes are known as proofreading mechanisms, in which the breaking of detailed balance enhances discrimination between correct and wrong substrates.

The first example is kinetic proofreading, introduced independently by Hopfield [3] and Ninio [4] in the 1970s. In this mechanism, the final readout does not occur immediately after enzyme– substrate binding, but only after a sequence of intermediate states in chemical space. Under nonequilibrium conditions, these states introduce a time delay between initial binding and product formation; since the complex with the wrong substrate typically has a larger dissociation rate, it has a higher probability of dissociating during this interval, thereby improving discrimination. Subsequent theoretical work has shown that proofreading is constrained by trade-offs between speed, accuracy and dissipation [5, 6, 7]. In classical kinetic proofreading, higher accuracy is typically obtained at the cost of slower selection, because repeated discrimination requires sufficiently long-lived intermediate states. This limitation has motivated alternative or complementary nonequilibrium strategies, such as catalytic discrimination, where the correct substrate promotes the catalytic or energy-consuming step more efficiently than the wrong one [8]. Such mechanisms can overcome the accuracy–speed trade-off and operate in combination with classical proofreading. Proofreading-like mechanisms have also been proposed in intracellular signaling, where multiple phosphorylation steps and phosphatase-mediated recycling can enhance specificity and reduce cross-talk between competing pathways [9, 10].

An alternative mechanism, known as spatial proofreading, was introduced in Ref. [11]. In this case, the selective delay does not arise from a sequence of intermediate chemical states, but from a sequence of different positions, i.e. from transport in physical space. The basic idea is to spatially separate the binding region from the readout region: during diffusive transport between these two regions, the wrong complex, being less stable, has a higher probability of dissociating before product formation. Of course, delayed stages should correspond to hard-driven transitions in order for proofreading to exceed the limits imposed by equilibrium thermodynamics. This can be realized for example through a sustained gradient of substrate. In this way, the spatial organization of the system, through concentration modulations and subcellular localization, can directly contribute to discrimination. This perspective makes spatial proofreading a potentially more general mechanism than classical kinetic proofreading, since it does not necessarily require the enzyme to have a sequence of intermediate states, nor that at least one step of the cycle be kept far from equilibrium through coupling to a chemical energy source. The nonequilibrium character is instead associated with steady spatial gradients that separate binding, transport and readout.

In the original work on spatial proofreading, specificity was shown to be controlled by the diffusive transport time, the substrate localization length scale and the association rate of the complex [11]. Moreover, a biologically plausible mechanism for generating substrate gradients was proposed, based on a phosphorylation–dephosphorylation cycle. This is consistent with the broader literature on cellular signaling, where gradients generated by localized kinase activity and distributed phosphatase activity are known to shape the propagation of phosphorylated species across the cell [12, 13, 14]. However, it would be desirable for spatial gradients to arise directly from nonequilibrium driving, namely from the chemical-potential imbalance associated with chemostatted nucleotide concentrations. In a thermodynamically consistent formulation, setting the nucleotide concentrations to their equilibrium values should remove these gradients and make the proofreading performance approach the equilibrium specificity (2). This requires microscopic reversibility and an explicit accounting of the chemical driving forces that maintain the system out of equilibrium [15, 16]. By contrast, in the original spatial proofreading framework [11], sustained substrate gradients were treated as imposed profiles, without explicitly introducing a microscopically reversible reaction network coupled to nucleotide chemostats and constrained by local detailed balance. As a consequence, the relation between specificity and thermodynamic driving is not explicitly determined by a microscopically reversible reaction network.

In this work, we introduce a thermodynamically consistent reaction–diffusion model of spatial proofreading. The gradient of active substrates is generated by a reversible phosphorylation– dephosphorylation cycle and the driving is explicitly controlled by the nonequilibrium chemical energy Δ*µ* associated with nucleotide chemostats,

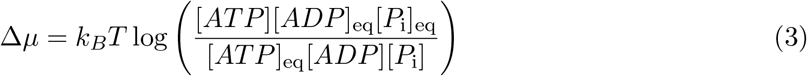

A further feature of the model is that the density of free proofreading enzyme is not imposed to be spatially uniform, but emerges self-consistently from the dynamics of the system. This aspect is relevant because enzyme availability can be limited in real biological contexts: complex formation can locally sequester a significant fraction of free enzyme, thereby modifying the effective binding kinetics and in turn discrimination.

The paper is organized as follows. We first analyze the model in a one-dimensional geometry, exploring its behavior as a function of the driving and of the kinetic parameters. We then extend the analysis to two-dimensional geometries, to study the role of spatial localization of the kinase-based substrate generation mechanism. Next, we turn to a stochastic description, to assess the effect of fluctuations in regimes of low molecular occupancy. Finally, we present a critical discussion of our results and possible extensions of our theoretical framework.

## 2. Results

### 2.1. A thermodynamically consistent model of spatial proofreading

We first introduce a thermodynamically consistent reaction–diffusion model where spatial proof-reading is coupled to a phosphorylation–dephosphorylation cycle driven out of equilibrium. A generic substrate *S* is converted into its active form *S*^*^ by a kinase localized at one end of the domain (e.g. a membrane), while *S*^*^ is converted back into *S* by a phosphatase distributed throughout the system. We assume that only activated substrates can bind to the enzyme. As a consequence, the spatial separation between localized activation and distributed deactivation generates a non-uniform profile of active substrates, providing the spatial structure required for the proofreading mechanism.

We first discuss our model in a one-dimensional geometry, which is sufficient for a preliminary analysis and allows us to highlight the fundamental physical mechanisms (see Fig. 1). The extension to two-dimensional domains will be discussed in a later section. We note that our model is fully general and can be readily adapted to arbitrarily designed compartmentalization schemes. In 1D the spatial domain is discretized into *n* compartments, labeled by *i* = 1, …, *n*, and we assume a uniform discretization step Δ*x*, so that the domain length is *L* = *n* Δ*x*.

**Figure 1.**
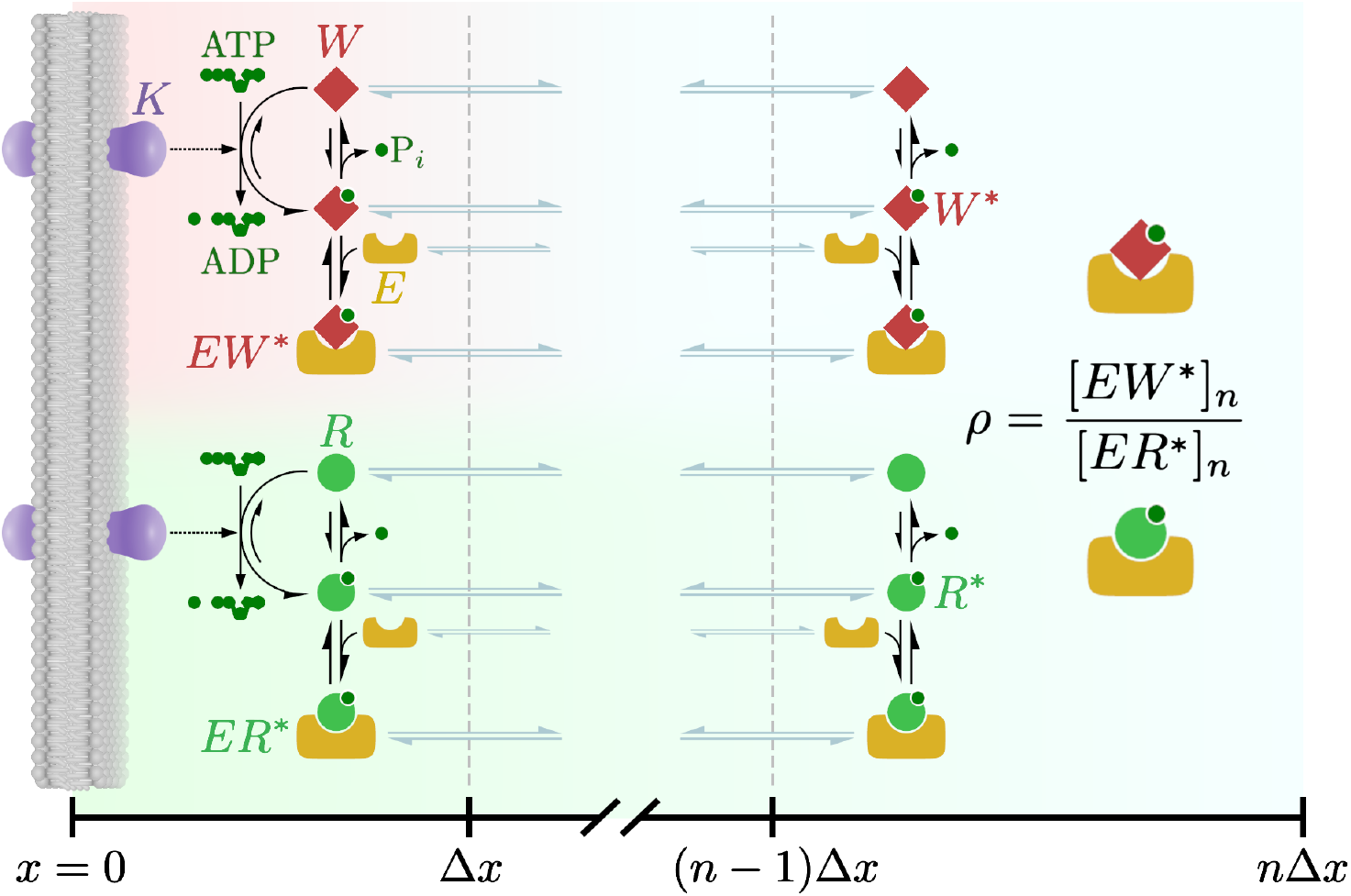
Schematic representation of the spatial proofreading model in one dimension. Substrate activation by a kinase occurs in the first compartment, while the phosphatase is distributed along the entire domain and catalyzes the deactivation of active species. Active substrates can bind to the enzyme, forming the complexes *ER*^*^ and *EW*^*^, which can dissociate with rates 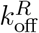 and 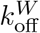. All dynamical species (except kinases) diffuse by jumping between adjacent compartments with rate *k*_*D*_ = *D/*Δ*x*^2^. The last compartment is identified as the readout region, where specificity is measured as *ρ* = [*EW*^*^]_*n*_*/*[*ER*^*^]_*n*_.

We consider two kinds of substrates, correct and wrong ones, denoted by *R* and *W*, respectively. The corresponding active forms, denoted by *R*^*^ and *W*^*^, are generated uniquely in the first compartment by kinase-catalyzed phosphorylation,

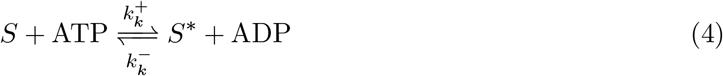

where *S* = (*R, W*). Concurrently, a phosphatase catalyzes deactivation (dephosphorylation) throughout the system (i.e. in all compartments),

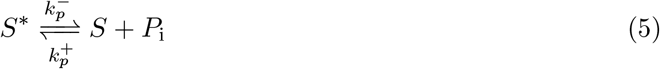

Here the *±* convention is chosen so as to identify the activation (*S → S*^*^) with the + direction. It should be noted that, since the kinase/phosphatase switch is modeled as two pseudo first-order reversible reactions, we effectively assume kinase and phosphatase activities to remain constant.

The proofreading enzyme *E* can bind to the active substrates, forming the complexes *ER*^*^ and *EW*^*^. As in kinetic proofreading, we assume that the only asymmetry between the correct and wrong substrates lies in the respective affinities of the complexes, identified here by the corresponding dissociation (off) rates. In particular, we have 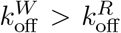. The wrong complex is less stable, and as a consequence it has a higher probability of dissociating during transport toward the readout region.

Thermodynamic consistency is imposed by requiring that, along the closed phosphorylation– dephosphorylation cycle, the ratio between the product of forward currents and that of backward currents is fixed by the thermodynamic driving force of the cycle,

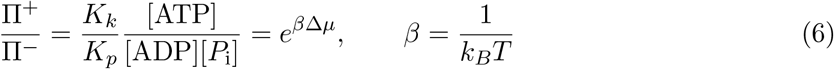

where 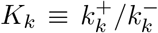 and 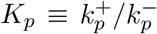 are the equilibrium constants of reactions (4) and (5), respectively. At equilibrium (Δ*µ* = 0), this condition reduces to detailed balance according to Kolmogorov’s criterion [17], thereby fixing the constraint

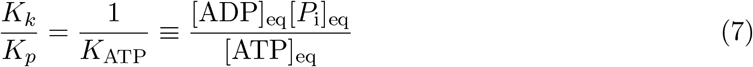

In the following, we choose *K*_*k*_ = 1, which gives *K*_*p*_ = *K*_ATP_. The details of the derivation and of the chemostat parametrization are reported in the Supplementary Materials.

Under nonequilibrium conditions, the driving is controlled by the concentrations of ATP, ADP and *P*_i_, treated as chemostats. In turn, these are computed as functions of Δ*µ* according to Eq. (3) (see Supplementary Materials). Therefore, in our model the strength of the nonequilibrium mechanism that sustains the gradient of active substrates can be tuned by varying the single parameter Δ*µ*. The typical driving corresponding to ordinary nucleotide concentrations in cells is Δ*µ* ≃20 *÷* 25 *k*_*B*_*T* [18].

Eq. (7) ensures that detailed balance is satisfied at equilibrium. Within this constraint, we are still free to adjust the *velocities* of the kinase and phosphatase steps. One simple way to do this is to introduce two kinetic scales, *v*_*k*_ and *v*_*p*_, to parametrize the phosphorylation (activation)– dephosphorylation (deactivation) rates. Overall, the local reactions considered in the model are reported in Tab. 1 along with the compartments where they occur.

In addition to chemical transformations, all species can diffuse between adjacent compartments. Assuming the same diffusion coefficient *D* for all species (we think of substrates as proteins of similar size to the proofreading enzyme), and well-mixed compartments, the spatial discretization fixes the jump rate *k*_*D*_ = *D/*Δ*x*^2^. We therefore add the reactions

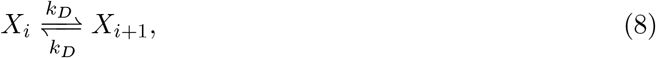

where *X*_*i*_ represents a generic chemical species *X* when it is localized in the *i −* th compartment. Reflecting boundary conditions are imposed – in the first and last compartments only jumps toward the interior of the domain are allowed. Since there are no sources nor sinks in our model, this ensures mass conservation. More precisely, for compartments of equal volume, the corresponding conservation laws can be written as

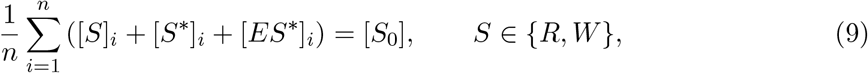

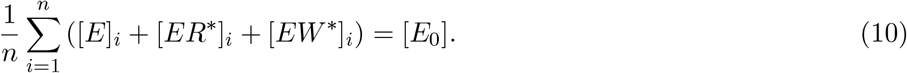

where [*S*_0_] and [*E*_0_] denote, respectively, the initial spatially uniform concentrations of substrate *S* and proofreading enzyme *E*.

Finally, our main observable is specificity as measured in the last compartment, identified as the readout region, that is

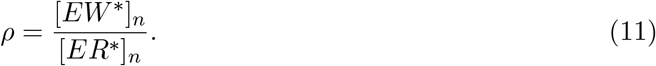

To quantify the improvement with respect to the equilibrium limit, we use the normalized specificity *ρ/ρ*_eq_. When association rates are equal and the initial concentrations of the two substrates are identical, [*R*_0_] = [*W*_0_], the equilibrium value is

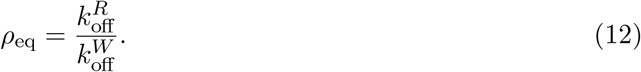

Values of *ρ/ρ*_eq_ below one indicate better discrimination than equilibrium, whereas values close to one correspond to the equilibrium limit.

As a first validation of the model, we analyze the dependence of the normalized specificity on the nonequilibrium driving *β*Δ*µ*, while keeping the other parameters fixed (Fig. 2a). As expected, for zero driving the system satisfies detailed balance, there is no substrate gradient (see Supplementary Materials) and the specificity approaches the equilibrium value. As *β*Δ*µ* increases, the phosphorylation–dephosphorylation cycle sustains a steeper and steeper gradient of active substrates and the spatial proofreading mechanism becomes effective. Consequently, the normalized specificity decreases, indicating improved discrimination with respect to the equilibrium limit. For sufficiently large values of the driving, the curve reaches a plateau. This indicates that, beyond a certain threshold, increasing driving strength is no longer the limiting factor for discrimination. In this regime, specificity is mainly controlled by the competition between diffusive transport, complex dissociation and deactivation of active substrates. A more detailed analysis of the response to driving, including the behavior of complex concentrations in the readout and parameter sweeps over the main kinetic and diffusive parameters is reported in the Supplementary Materials.

**Figure 2.**
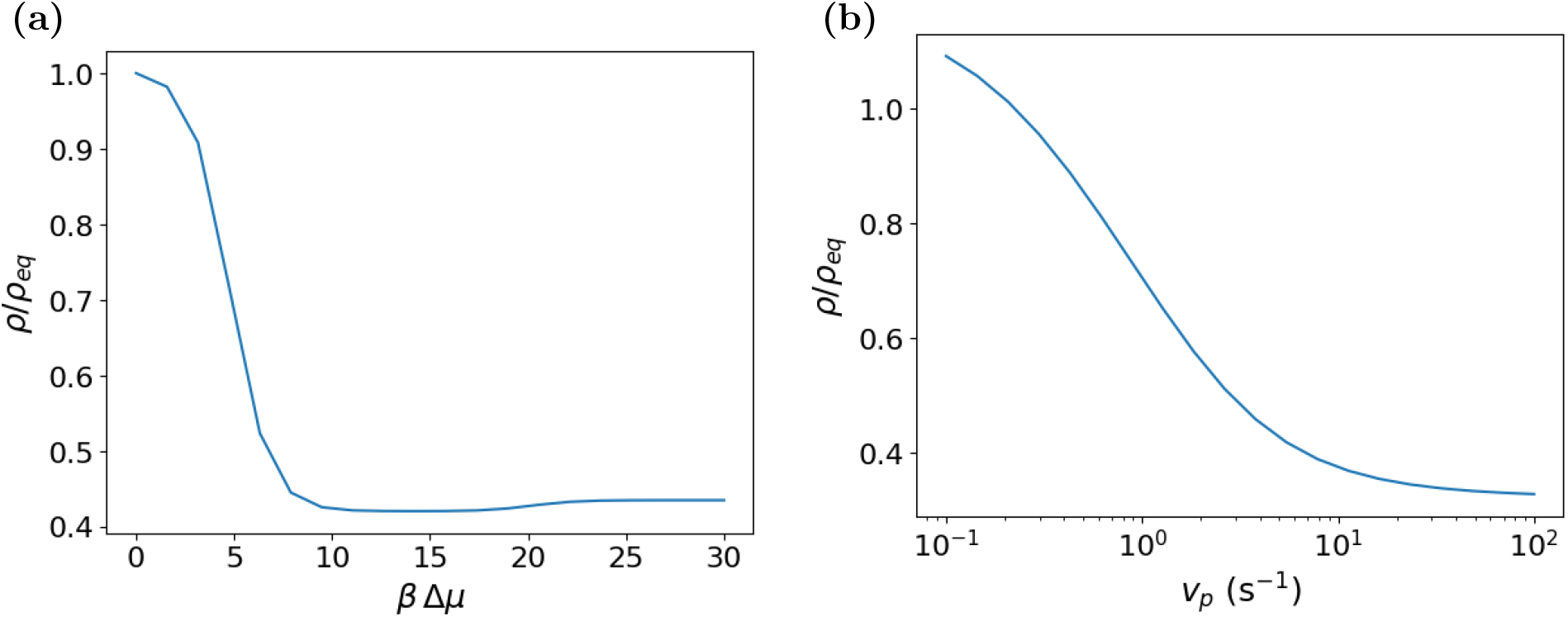
Normalized specificity *ρ/ρ*_eq_, computed from Eq. (11) in the steady state, as a function of the nonequilibrium driving *β*Δ*µ* **(a)** and of the phosphatase rate *v*_*p*_ **(b).** Unless stated otherwise, the reference parameter set is: *β*Δ*µ* = 20, *n* = 10, Δ*x* = 1 *µ*m (*L* = 10 *µ*m), *D* = 10 *µ*m^2^ s^*−*1^, *k*_on_ = 10^6^ M^*−*1^s^*−*1^, 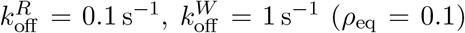, *v*_*k*_ = 3.0 *×* 10^2^ M^*−*1^s^*−*1^, *v*_*p*_ = 5 s^*−*1^, *K*_*k*_ = 1, *K*_*p*_ = *K*_ATP_ = 3.9 *×* 10^*−*7^ M^*−*1^, [*R*_0_] = [*W*_0_] = 3.0 *µ*M, and [*E*_0_] = 0.6 *µ*M. A detailed discussion of the reference parameter choice is provided in the Supplementary Materials.

### 2.2 One-dimensional parameter regimes

In this section, we identify the key parameters governing specificity in the one-dimensional domain. Our goal is not only to describe how *ρ/ρ*_eq_ depends on each parameter, but also to pinpoint the physical nonequilibrium mechanisms that control spatial proofreading.

In the original work, the role of rebinding was gauged by the effective binding rate *k*_*b*_ = *k*_on_*ρ*_*E*_, where *ρ*_*E*_ denotes the density of free enzyme, which was assumed to be constant along the domain [11]. By contrast, in our model the distribution of free enzyme emerges self-consistently from the coupled dynamics of binding, dissociation and diffusion. Therefore, the effects of *k*_on_ and total enzyme concentration *E*_0_ can be analyzed separately within our theoretical framework.

The results obtained are consistent with the results reported in Ref. [11] – an increase in the effective probability of rebinding tends to reduce the discriminatory ability of the system (see Supplementary Materials). In the following sections, we focus on the parameters that more directly control substrate gradient as the driving force of discrimination, namely diffusive transport and selective complex dissociation.

#### The phosphatase controls the confinement of the active-substrate gradient

To quantify the role of phosphatase-controlled deactivation, we performed a logarithmic sweep of the parameter *v*_*p*_, which controls the deactivation rate of active substrates (see Table 1). Figure 2b shows that increasing *v*_*p*_ significantly reduces *ρ/ρ*_eq_, until an asymptotic plateau is reached. This behavior reflects the role of the phosphatase in spatially confining the gradient profile of active substrates. Faster deactivation reduces the characteristic length 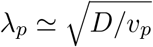 over which active species can diffuse before being converted back into the inactive form.

**Table 1.** Local reactions considered in the model and corresponding rates, with explicit indications of the compartment where they are present. The quantities *k*^+^ and *k*^*−*^ indicate the forward (activation, left-to-right) and backward (deactivation) rates, respectively. Note that *k*^*±*^ for the kinase, as well as *k*^+^ for the phosphatase, are second-order rates. However, the corresponding reactions become pseudo first-order in our scheme, as the concentrations of nucleotides and inorganic phosphate are chemostatted as specified by the driving Δ*µ*.

| Reaction | | $k^+$ | $k^-$ | Domain |
| --- | --- | --- | --- | --- |
| Kinase | $R + \text{ATP} \rightleftharpoons R^* + \text{ADP}$ | $K_k v_k$ | $v_k$ | $i = 1$ |
| | $W + \text{ATP} \rightleftharpoons W^* + \text{ADP}$ | $K_k v_k$ | $v_k$ | $i = 1$ |
| Phosphatase | $R + P_i \rightleftharpoons R^*$ | $K_p v_p$ | $v_p$ | $i = 1, \dots, n$ |
| | $W + P_i \rightleftharpoons W^*$ | $K_p v_p$ | $v_p$ | $i = 1, \dots, n$ |
| Complex formation | $E + R^* \rightleftharpoons ER^*$ | $k_{\text{on}}$ | $k_{\text{off}}^R$ | $i = 1, \dots, n$ |
| | $E + W^* \rightleftharpoons EW^*$ | $k_{\text{on}}$ | $k_{\text{off}}^W$ | $i = 1, \dots, n$ |

From the proofreading point of view, for fixed values of the other parameters, increasing *v*_*p*_ makes the removal of active substrates in the readout region more efficient. However, this mainly penalizes the wrong substrate: the complex *EW*^*^, having a shorter lifetime, dissociates more frequently releasing *W*^*^, which, once free, is exposed to phosphatase-catalyzed deactivation. By contrast, the correct complex *ER*^*^ remains bound for a longer time on average and is therefore partially protected from this process. The result is a reduction of the ratio [*EW*^*^]_*n*_*/*[*ER*^*^]_*n*_ and hence an improvement in discrimination.

In the opposite regime, when *v*_*p*_ is small, dephosphorylation is slow compared with diffusive transport. For this reason, active substrates can diffuse over longer distances before being converted back into the inactive form, making the gradient less sharp. The readout region is no longer sufficiently separated from the activation source and the spatial mechanism becomes less effective: the wrong substrate is not selectively penalized and *ρ/ρ*_eq_ can approach unity or even exceed it, indicating discrimination comparable to, or even worse than, the equilibrium limit.

An analogous control was performed by varying the catalytic activation rate of the kinase, *v*_*k*_. We found that specificity is only weakly affected by this parameter: changing *v*_*k*_ mainly modifies the flux of active substrates injected into the domain, without significantly altering the confinement mechanism controlled by the phosphatase. The details of this analysis are reported in the Supplementary Materials.

#### Diffusion generates an optimal discrimination window

We now turn to analyze the role of diffusion on spatial proofreading. The key parameter is the characteristic transport time across the domain, *τ*_*D*_ = *L*^2^*/D*. This time sets the delay required to achieve proofreading, i.e. the duration of the time window during which complexes can dissociate before reaching the readout region. For this reason, diffusion affects the effectiveness of differential dissociation between the correct and wrong complexes. Figure 3 shows the normalized specificity *ρ/ρ*_eq_ as a function of *τ*_*D*_ for different combinations of dissociation rates 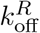 and 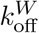. Interestingly, the curves do not collapse when the ratio 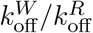 is kept fixed. Therefore, discrimination does not depend only on the relative magnitude of the two dissociation rates, but also on their absolute values, a clear blueprint of nonequilibrium.

**Figure 3.**
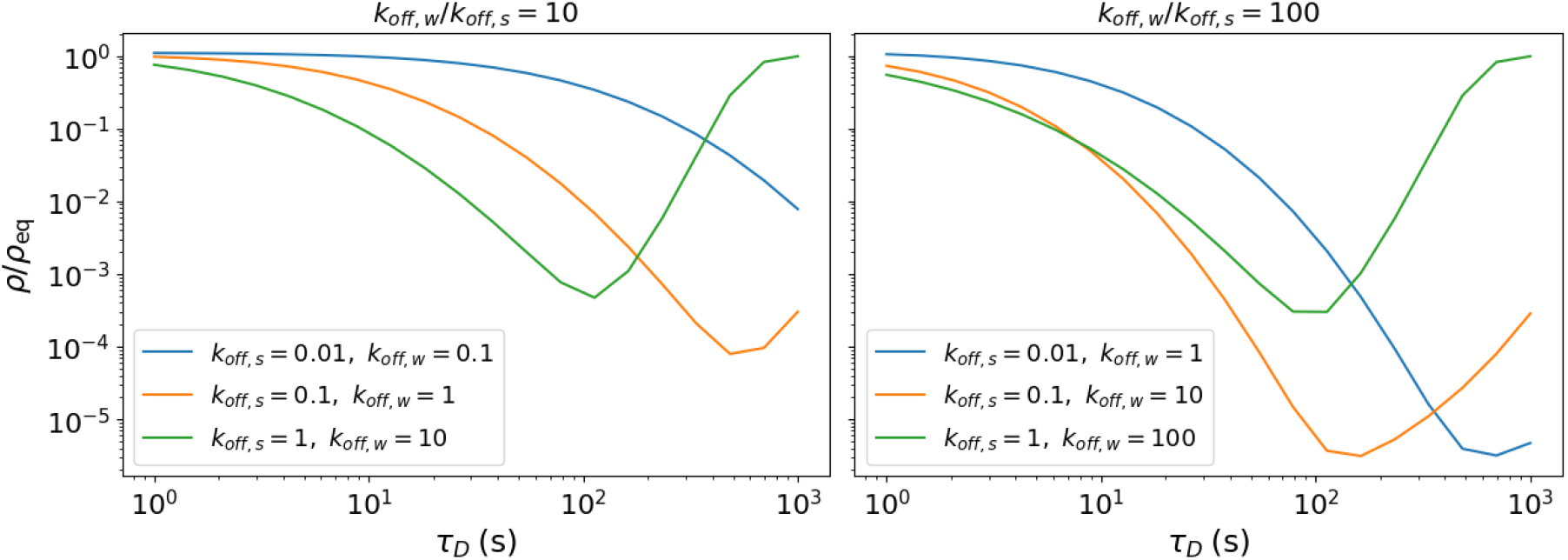
Normalized specificity *ρ/ρ*_eq_ as a function of the diffusive transport time *τ*_*D*_ = *L*^2^*/D* for different combinations of dissociation rates. The diffusion coefficient *D* was varied logarithmically in the range [10^*−*1^, 10^2^] *µ*m^2^ s^*−*1^ at fixed *L* = 10 *µ*m. A more detailed analysis reported in the Supplementary Materials shows that *τ*_*D*_ is the relevant scaling variable. The remaining parameters are those used in Fig. 2.

For small values of *τ*_*D*_ (fast diffusion limit), the normalized specificity approaches values close to unity. In this regime, transport is fast compared with dissociation times and concentration profiles tend to become uniform along the domain. Hence, a spatial separation between the activation and the readout regions has little effect, and the system approaches the equilibrium limit in a well-mixed environment. As *τ*_*D*_ increases, the wrong complex has a longer time window to dissociate before reaching the readout region. Consequently, *ρ/ρ*_eq_ decreases and discrimination improves. However, as transport becomes too slow, active substrates and complexes start localizing closer and closer to the source region, save for the same small local contribution in each compartment due to the backward phosphatase-catalyzed reaction. Eventually, as *τ*_*D*_ → ∞, transport is lost altogether, stationary proofreading happens almost exclusively in the first compartment and the specificity therefore approaches equilibrium. As a result, spatial proofreading is not monotonic: the normalized specificity displays a minimum at intermediate values of *τ*_*D*_. At fixed affinity ratio, increasing the absolute values of 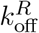 and 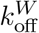 shifts the region of maximal discrimination toward shorter *τ*_*D*_. This behavior suggests that the optimal region corresponds to locking between transport and dissociation time scales of the complexes.

The existence of an optimal region indicates that spatial proofreading requires a compromise – transport must be slow enough to allow selective dissociation of the wrong complex, but not so slow as to almost completely isolate the readout region from the activation source. When *τ*_*D*_ is too long, diffusive coupling across the domain becomes weak and the dynamics in each compartment is dominated by local kinetics. We emphasize that, in the limit *D* → 0, the readout region becomes effectively decoupled from the kinase-driven source. Since the model is thermodynamically consistent, active substrates can then be generated locally only through the reverse phosphatase reaction, while the influence of the nonequilibrium driving becomes negligible at the readout. As a result, the specificity tends toward its equilibrium value.

To clarify further the origin of the minimum, we analyze separately the concentrations of the complexes in the last compartment (Fig. 4a) and the local contributions to the dynamics of *EW*^*^ (Fig. 4b) for the reference parameter set. As *τ*_*D*_ increases, the concentrations of both complexes decrease in the readout region, but beyond the optimal value *EW*^*^ tends to reach a plateau, whereas *ER*^*^ keeps decreasing. As a consequence, the ratio [*EW*^*^]_*n*_*/*[*ER*^*^]_*n*_ increases and discrimination worsens. Moreover, in this regime the absolute concentrations of the complexes become very low everywhere away from the source (reducing to the residual phosphorylation catalyzed by the phosphatase). Incidentally, this makes fluctuations associated with finite molecule numbers relevant, as we show later in the paper.A dedicated analysis of the trade-off between discrimination and the correct-complex concentration at the readout is reported in the Supplementary Materials, including these low-concentration regimes.

**Figure 4:**
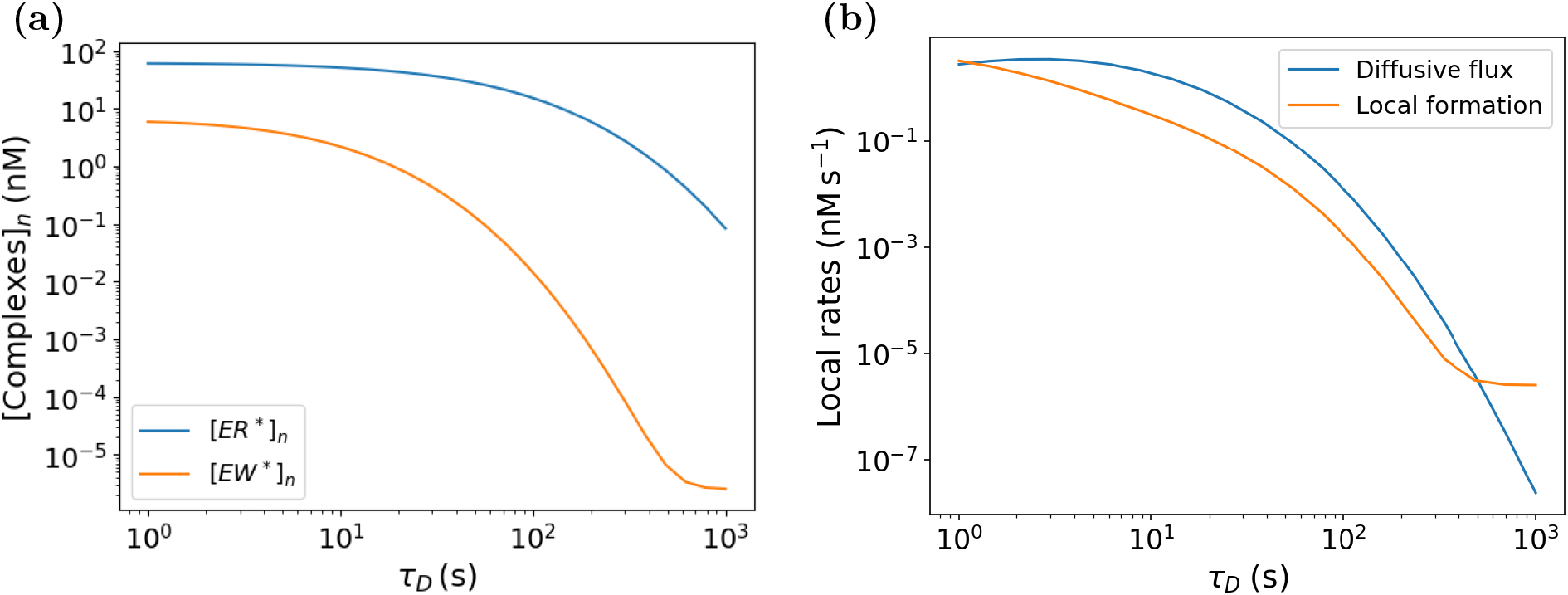
**(a)** Steady-state concentrations of the complexes [*ER*^*^]_*n*_ and [*EW*^*^]_*n*_ in the last compartment. **(b)** Comparison between the incoming diffusive contribution |*J*_*n−*1*→n*_|*/*Δ*x* and the local formation term 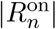 for *EW*^*^. The parameters are the same as those used in Fig. 2.

The local balance in the last compartment can be written as

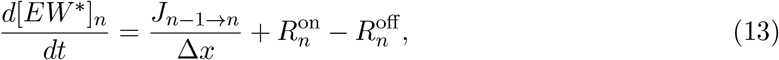

where *J*_*n−*1*→n*_ is the incoming diffusive flux, 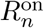 is the local formation term of the complex, and *R*^off^ the dissociation term. The change of regime occurs when the transport contribution becomes comparable to the local formation term (see also the orange curve in Fig. 3a). In this limit, further reducing *D* no longer enhances the selective penalization of the wrong complex during transport. Instead, it only reduces the diffusive supply of active species toward the readout region.

The role of diffusion described above is consistent with the interpretation proposed in Ref. [11]. In the presence of a gradient, transport between the binding and readout regions provides the delay required to generate a strongly driven sequence of testing stages for the wrong substrate. These are the essential ingredients of proofreading. In our model, however, the steady gradient of active species is not imposed externally, but is self-consistently generated by a reversible phosphorylation–dephosphorylation cycle driven far from equilibrium. The minimum observed as a function of diffusion therefore reflects not only the balance between transport and dissociation times, but also the local balance between diffusive supply and complex formation in the readout compartment.

A limitation of the present description is the assumption of the same diffusion coefficient for all species. In a biological context, enzyme–substrate complexes may diffuse more slowly than free species because of their larger size. Additional analyses (see Supplementary Materials) showed that improving discrimination does not necessarily require a global reduction of the diffusivity of all species: it may be sufficient that the complexes *ER*^*^ and *EW*^*^ diffuse more slowly, thereby increasing the time window available for selective dissociation during transport down the gradient.

#### Dissociation rates define the temporal window for discrimination

Selective dissociation provides the very definition of right and wrong substrates. It is therefore necessary to investigate how spatial proofreading depends on the corresponding rates, 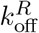 and 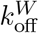. These parameters must be considered as distinct, since discrimination does not depend only on their ratio, but also on their absolute values. Figure 5a shows that the best discrimination is obtained in an intermediate region of parameter space, where 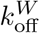 is large enough to allow dissociation of the wrong complex during transport, while 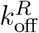 remains small enough to allow the correct complex to reach the readout region before dissociating. By contrast, along the diagonal 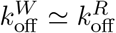, the two complexes have similar stability and the spatial mechanism cannot amplify discrimination, so that *ρ/ρ*_eq_ remains close to unity. To distinguish the effect of the ratio between the dissociation rates from that of their absolute values, we fix the ratio 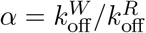 and let 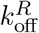 vary. High values of *α* correspond to very unstable wrong complexes with respect to the right ones. As shown in Fig. 5b,for *α >* 1 the normalized specificity displays a minimum at intermediate values of 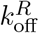. This confirms that increasing the wrong-to-right affinity ratio is not sufficient: proofreading is effective only when dissociation times are comparable with the transport time. The natural parameter that quantifies this balance between different physical time scales is the Damköhler numbers. For the two complexes we define

**Figure 5:**
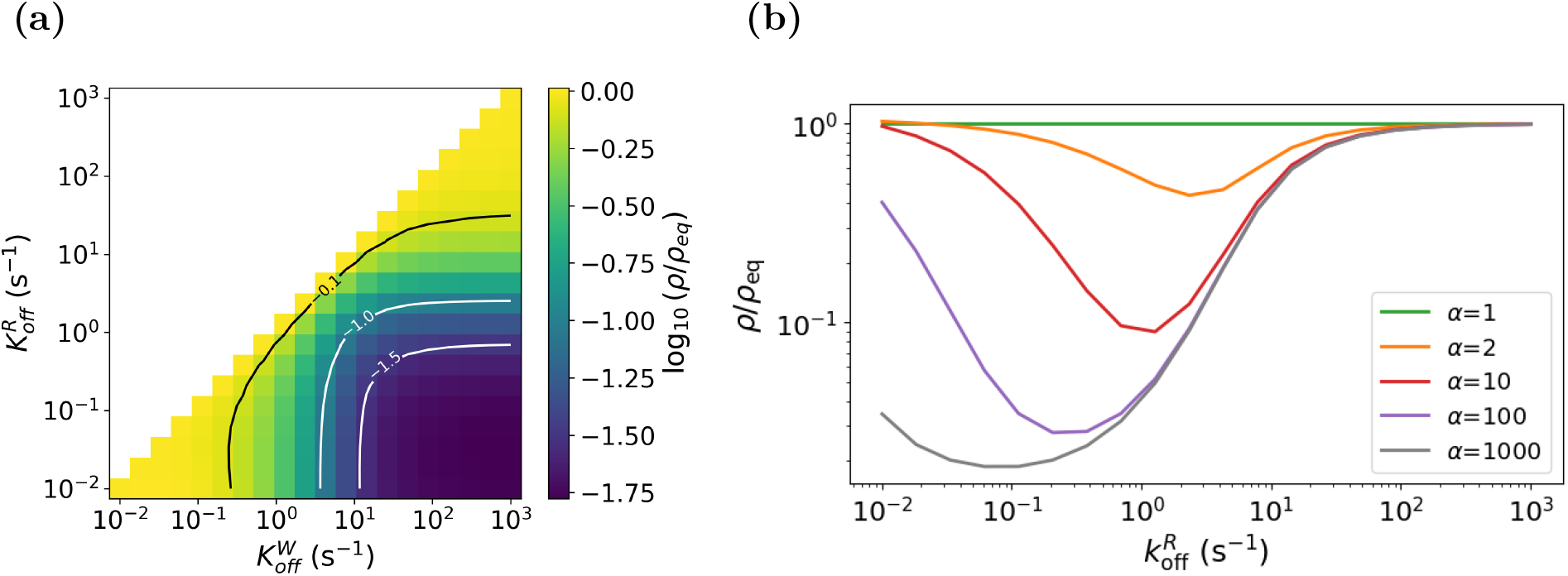
**(a)** Heatmap of normalized specificity in the half-plane 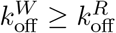. **(b)** Normalized specificity as a function of 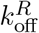for different values of the ratio 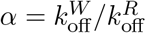. Other parameters as in Fig. 2.

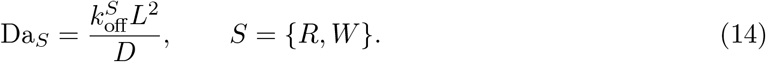

These parameters gauge the characteristic diffusive transport time across the domain, *τ*_*D*_ ~ *L*^2^*/D* = 10 s, relative to the mean lifetimes of the complexes, 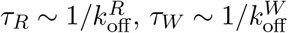.

For Da_*R*_, Da_*W*_ ≪ 1, dissociation is slow compared with transport: both complexes survive long enough to reach the readout region and proofreading is weak. In the opposite regime, Da_*R*_, Da_*W*_ ≫ 1, even the correct complex dissociates too rapidly, reducing the amount of *ER*^*^ available in the readout region. Hence, optimal spatial proofreading emerges in an intermediate window, where the wrong complex is penalized during transport, Da_*W*_ ≫ 1, while the correct one retains a significant probability of reaching the final region, i.e. Da_*R*_ ≃ 1. That is to say, optimal discrimination in a reaction-diffusion context requires kinetic control of the chemical steps involving wrong substrates and a kinetic/diffusive balance for those involving right substrates.

### 2.3. In two dimensions, source localization shapes specificity at the readout boundary

Our thermodynamically consistent one-dimensional model of spatial proofreading captures the essential physics of error reduction through nonequilibrium control of spatially separated testing stages. Yet, by construction, it cannot resolve the role of source and readout geometry. We therefore extend the analysis to a two-dimensional domain, allowing us to assess how spatially localized kinases modulate specificity along the readout boundary. To this end, we consider a two-dimensional grid, which allows us to introduce a transverse direction with respect to the main transport axis. In this geometry, substrate activation is not necessarily uniform along the source boundary, but can be confined to specific regions. This makes it possible to study whether and how discrimination measured on the opposite boundary is affected by the geometry of the activation source.

Specifically, we consider a rectangular domain of size *W× H*, discretized with spatial steps Δ*x* = Δ*y*. Kinase activity is localized on the *x* = 0 (source) boundary, while specificity is measured along the *x* = *W* (readout) boundary, both extending along the *y* direction. Diffusion is implemented with reflecting boundary conditions along *x* and periodic boundary conditions along *y*. To isolate the effect of geometry, we compare two representative configurations at fixed total catalytic area, that is, with the same number of cells where kinase activity is present – a distribution of equally spaced bands and a single central band. The local value of *v*_*k*_ is adjusted to keep the overall source strength constant ^1^, so that any differences in specificity can be attributed to the spatial distribution of the activation source and not to a different global intensity of the activation process. Figure 6 shows that the spatial distribution of kinases is reflected in the profile of the normalized specificity along the readout boundary. In the case of equally spaced bands, *ρ*(*y*)*/ρ*_eq_ displays only moderate oscillations around the value obtained for a uniform kinase distribution of the same strength. This indicates that the profiles generated by the different activation regions overlap through lateral diffusion: each point of the readout boundary receives contributions from multiple sources and specificity is averaged along the transverse direction. In the case of a single central band, instead, the profile is more heterogeneous: discrimination is reduced in the region of the readout aligned with the kinase band, where *ρ*(*y*)*/ρ*_eq_ takes larger values. In this region, active species reach the right boundary through more direct diffusive paths and therefore with shorter transport times. Moving toward the lateral regions, the effective transport time increases, because the species must also diffuse along the transverse direction. This makes the proofreading mechanism more effective away from the central region: the wrong complex *EW*^*^ has a higher probability of dissociating during transport and the wrong active substrate *W*^*^, once free, has more time to be deactivated by the phosphatase. Consequently, the contribution of wrong species to the readout decreases and specificity improves. The same marked decrease in *ρ*(*y*)*/ρ*_eq_ is not observed in the case of equally spaced bands, because the presence of multiple sources distributed along *y* favors the lateral overlap of the activation profiles. In this case, the effect of different diffusive times is partially averaged out and the specificity profile remains more uniform. Further comparisons between distributed and centralized kinase patterns, together with additional spatial concentration maps and an analysis of the effect of diffusion, are reported in the Supplementary Materials.

**Figure 6.**
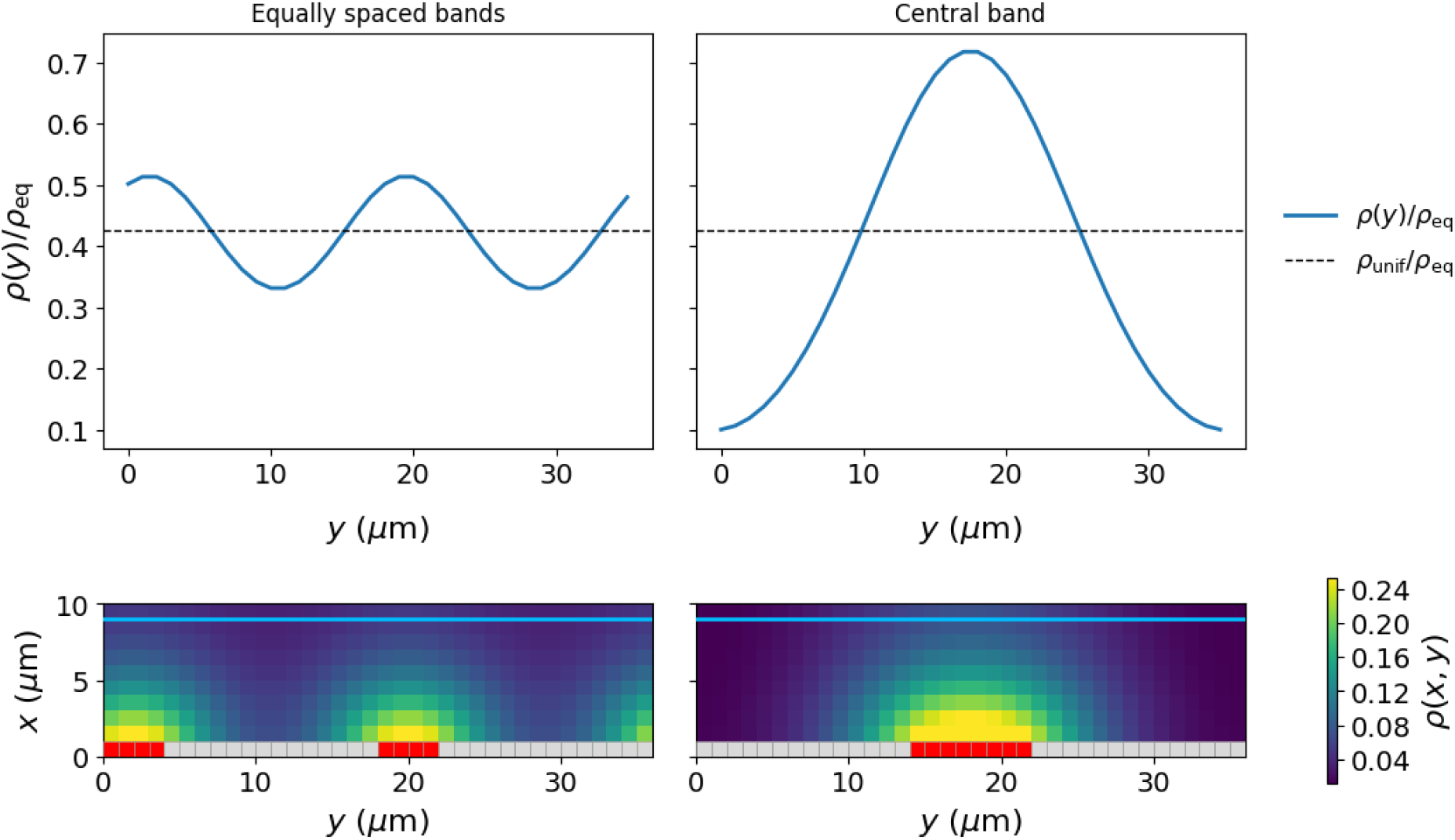
Normalized specificity profiles along the readout boundary for different equal-strength source geometries. Left panels: equally spaced band distribution of sources, with bands of width *L*_*y*_ = 4 *µ*m separated by a distance *d* = 14 *µ*m. Right panels: single source band of width *L*_CB_ = 8 *µ*m at the center. The horizontal dashed lines in the upper plots indicate the normalized reference value obtained for a uniform kinase distribution of the same strength, *ρ*_unif_ */ρ*_eq_≃ 0.43. The lower panels show the corresponding two-dimensional maps of the local specificity. In the first compartment, red/grey cells indicate the presence/absence of kinases. The cyan line separates the last compartment, *x* = *W*, identified as the readout region. The domain has size *W* = 10 *µ*m and *H* = 36 *µ*m, with discretization Δ*x* = Δ*y* = 1 *µ*m. Other parameters as in Fig. 2.

To assess to what extent the spatial modulation of specificity depends on the geometry of the domain, we also considered a circular two-dimensional geometry. In this case, the domain consists of a circular cellular region with a central nucleus excluded from diffusion. The kinase is localized on the external boundary of the nucleus, while specificity is measured along the internal membrane boundary of the cell. This geometry could model e.g. perinuclear kinase-controlled trafficking from the trans-Golgi network, where localized kinase activity, for example by protein kinase D, phosphorylates components of the export machinery and thereby regulates the delivery of activated cargoes to distant plasma-membrane readout sites (although in this case active transport should probably be factored in the model). The readout profile is parametrized by the angle *θ*, defining

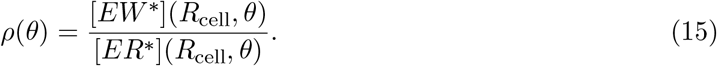

Figure 7 shows that, also in the circular domain, kinase localization generates a spatial modulation of specificity along the readout boundary. When kinases are distributed symmetrically around the nucleus, the normalized profile *ρ*(*θ*)*/ρ*_eq_ remains close to the mean value of the corresponding case in which kinases are uniformly distributed over the entire peri-nuclear region^2^. This indicates that the contributions from the different activation regions overlap significantly and produce a relatively uniform readout. By contrast, when kinase activity is confined to a single band, specificity shows a marked angular dependence. In particular, the ability to discriminate is reduced in the region of the external boundary directly facing the source, while it improves substantially in the more distant angular regions behind the source. This behavior is consistent with the physical mechanism emerging from the rectangular domain. Points of the readout reached through more direct diffusive paths receive a larger contribution from wrong species that are still active, whereas longer diffusive paths favor dissociation of the wrong complex and the subsequent deactivation of the free wrong substrate. Overall, these results suggest that specificity is governed not only by local reaction kinetics, but also by how the relative source–readout geometry shapes the diffusive paths of the complexes.

**Figure 7.**
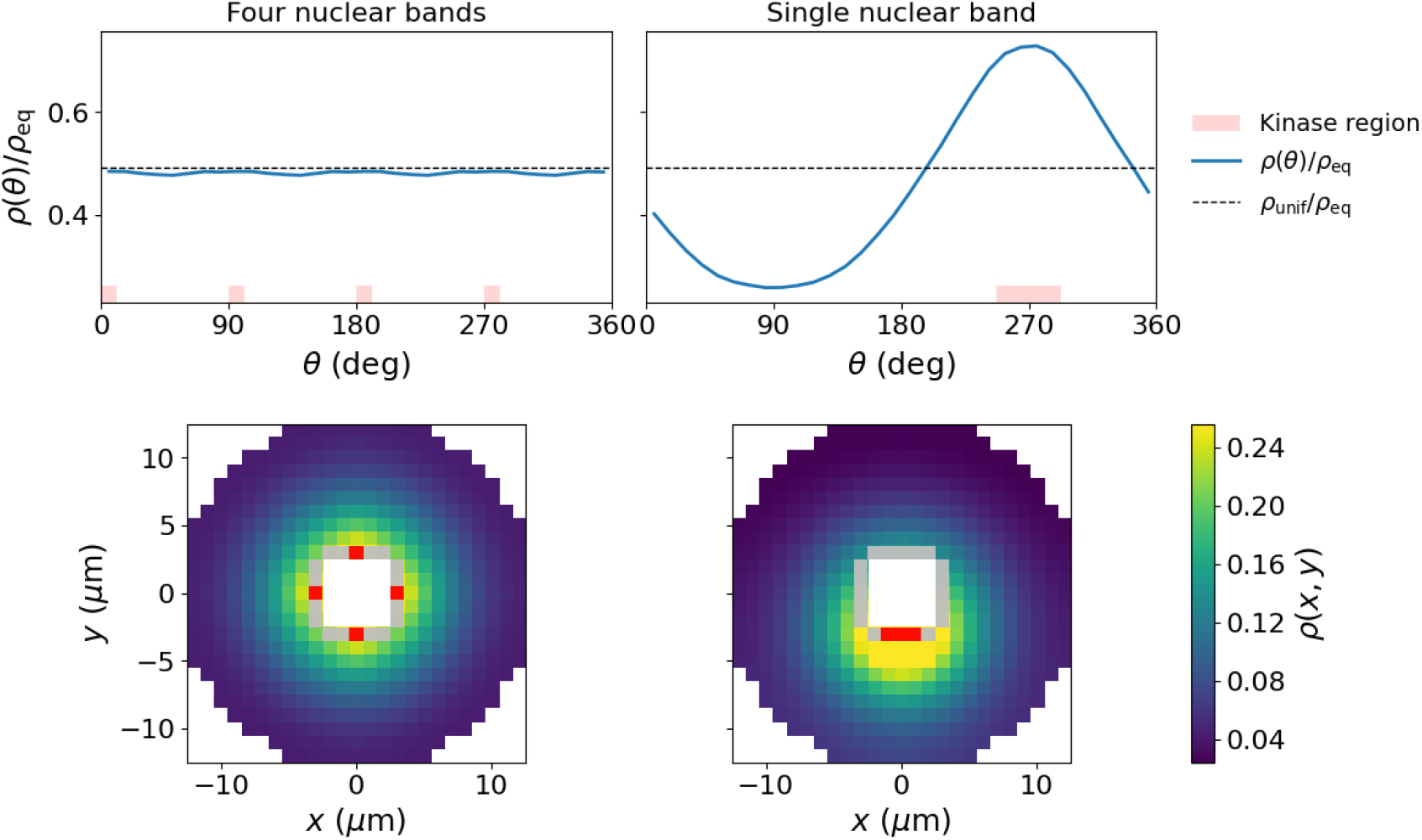
Normalized specificity profiles along the outer circular readout boundary for different equal-strength peri-nuclear source geometries (red regions at the bottom of the top panels). Left panels: symmetric kinase distribution over four localized regions on the sides of the nucleus. Right panels: a single kinase band localized on one side of the nucleus. In the upper panels, the dashed line corresponds to the uniform peri-nuclear kinase distribution, with *ρ*_unif_ */ρ*_eq_≃ 0.49. The lower panels show the two-dimensional maps of the local unnormalized specificity *ρ*(*x, y*). Red/grey cells indicate the presence/absence of a source (kinase). The domain has cellular radius *R*_cell_ = 13 *µ*m, nuclear radius *R*_nuc_ = 3 *µ*m and discretization Δ*x* = Δ*y* = 1 *µ*m. Other parameters as in Fig. 2.

### 2.4. Stochastic simulations reveal intermittent fluctuations in specificity

The previous analyses were performed using deterministic simulations, where concentrations represent the average behavior within well-stirred compartments with large copy numbers. However, in many biological contexts, the biomolecules involved in signaling and discrimination processes can be present in low copy numbers. As a consequence, intrinsic fluctuations associated with the discrete nature of reaction events may become relevant, especially for low-occupancy species such as the complexes in compartments far from the substrate sources. To assess this effect, we simulated the stochastic dynamics of the model using the Gillespie algorithm [19]. In stochastic regimes, proofreading performance cannot always be characterized only by steady-state mean concentrations, since the relevant readout may depend on rare events, first-passage times, threshold crossings, or product-counting strategies [20].

As a first control, we compare the time evolution of the complexes *ER*^*^ and *EW*^*^ in the last compartment of the 1D geometry with the deterministic prediction obtained with the same parameters. After a short initial transient, the stochastic trajectories fluctuate around the deterministic steady-state values, as shown in Fig. 8a. This behavior confirms that the deterministic dynamics correctly describes the mean value of the system, while the stochastic approach allows us to characterize temporal fluctuations and the intermittency of binding events at low copy numbers of complexes. To a closer look, however, fluctuations reveal a more subtle pattern (Fig. 8b). The distribution of stochastic specificity seems to display two populations, both revealing better discrimination than equilibrium, but one of them outperforming the proofreading ability of the deterministic prediction with the same set of parameters. This seems to indicate that fluctuations can in some cases boost spatial proofreading.

**Figure 8:**
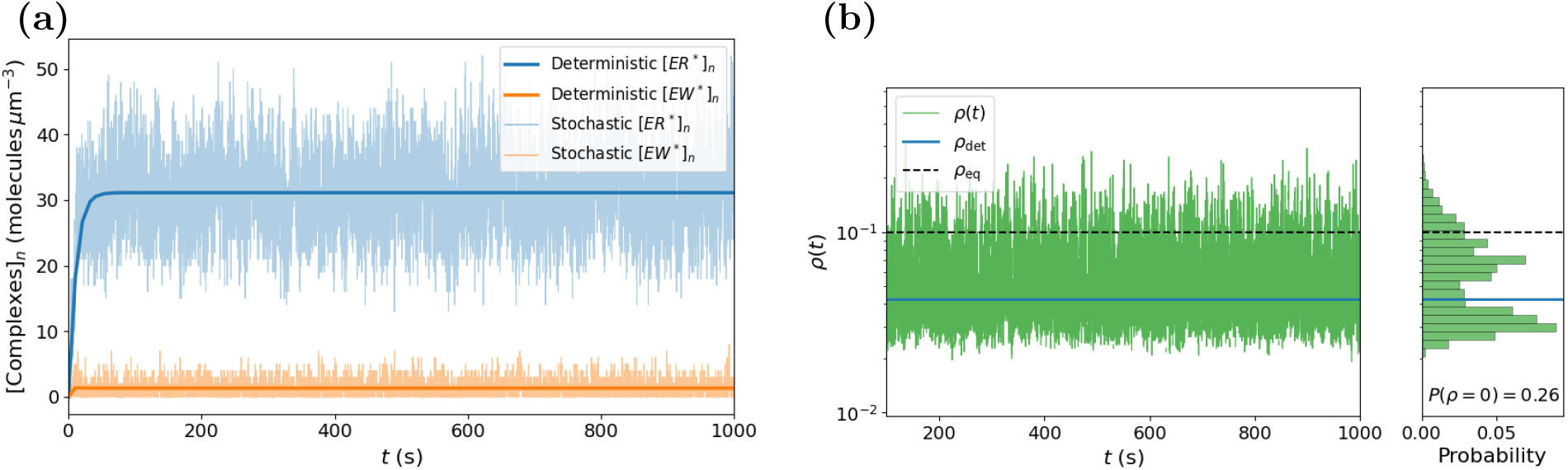
**(a)** Time evolution of the population of correct and wrong complexes, *ER*^*^ and *EW*^*^, in the last compartment of the [0, *L*] 1D geometry simulated with the Gillespie algorithm. The stochastic trajectories show fluctuations around the deterministic prediction (thick solid lines). **(b)** Time series of the instantaneous stochastic specificity, computed after the initial transient *t*_0_ = 10^2^ s (left) and probability distribution of the strictly positive values of *ρ*(*t*) (right). Solid line: deterministic value *ρ*≃ _det_ 4.27 *×* 10^*−*2^. Dashed line: equilibrium value *ρ*_eq_ = 0.1. Time points with *ρ*(*t*) = 0 (not reported) amount to *P* (*ρ* = 0) = 0.26. Other parameters are as in Fig. 2.

Even more interesting, about 25% of the time, stochastic discrimination appears to be *exact*, simply meaning that for about 25% of the time there are strictly no wrong complexes at the readout boundary. This is obviously an occurrence that cannot be observed at the mean-field level. Indeed, in the stochastic regime the wrong complex *EW*^*^ can be present intermittently – long intervals without any wrong complex can be interrupted by short episodes where *EW*^*^ appears in the readout compartment. The correct stochastic metric must therefore account for the relative amplitude of the error when both complexes are present, but also for the probability that a wrong episode indeed occurs. To this end, we define an event-weighted, strictly positive stochastic specificity,

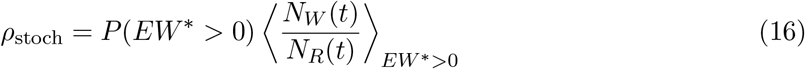

where *N*_*W*_ (*t*) and *N*_*R*_(*t*) denote, respectively, the number of *EW*^*^ and *ER*^*^ complexes in the read-out compartment, and the conditional average is computed only over time intervals during which at least one wrong complex is present. This definition explicitly separates the two contributions: the probability of observing a wrong event, *P* (*EW*^*^ *>* 0), and the mean ratio between wrong and correct complexes during such episodes. In the limit in which *EW*^*^ is persistently present and fluctuations are small, this definition reduces to the time average of the ratio *N*_*W*_ */N*_*R*_, which approaches the deterministic description. In the reference parameter regime, after excluding the initial transient, we obtain

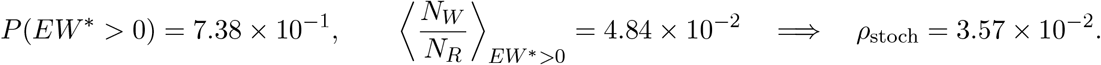

The dispersion of wrong episodes was estimated from the standard deviation of the ratio *N*_*W*_ */N*_*R*_ computed over the intervals in which *EW*^*^ *>* 0. The stochastic specificity is obtained by multiplying the conditional ratio by the probability of observing a wrong event. Therefore, the effective dispersion associated with *ρ*_stoch_ is estimated as 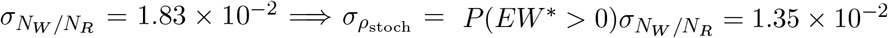. Thus,

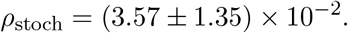

Within the episode-to-episode dispersion, this value is compatible with the deterministic prediction *ρ*_det_ ≃ 4.27 *×* 10^*−*2^. However, the stochastic description shows that the wrong readout is not determined by a continuous concentration, but by intermittent episodes of *EW*^*^ in the readout compartment. In the case considered, *EW*^*^ is present for a significant fraction of the time and appears in distinct episodes with mean duration ≃ 0.37 s. In this sense, this description makes it possible to distinguish between weak and strong errors characterized by different temporal patterns (see again Fig. 8b).

The event-weighted definition of specificity provides a sensible measure in the stochastic regime, particularly when the probability of finding the wrong complex in the readout region is strongly suppressed. This occurs, for example, for a low diffusion coefficient *D* ≃ 0.5 *µ*m^2^ s^*−*1^, as shown in

Fig. 4a. To put this value in perspective, this diffusion coefficient is slow for a freely diffusing cytosolic protein, but lies within the range reported for the lateral diffusion of membrane-associated or transmembrane proteins in crowded eukaryotic membranes and is comparable to the faster diffusive component reported for cytoplasmic mRNA/mRNP particles [18]. In this regime, we obtain

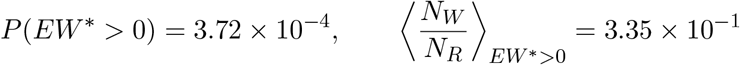

The first value shows that the wrong complex is present in the readout compartment only for a very small fraction of the time, while the second indicates that, when a wrong event occurs, the instantaneous ratio between wrong and correct complexes can be non-negligible. The corresponding stochastic specificity is

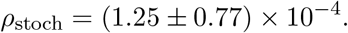

For the same parameters, the deterministic description gives *ρ*_det_ = 1.21 *×* 10^*−*4^, in agreement with the stochastic value within the dispersion associated with the wrong episodes. Consequently, the new definition provides additional information on the nature of errors in the stochastic, low-diffusion regime: the low value of *ρ*_stoch_ does not arise because the ratio *EW*^*^*/ER*^*^ is always uniformly small in time, but because the episodes in which the wrong complex reaches the readout are extremely rare. The metric *ρ*_stoch_ is therefore particularly useful in rare-event regimes, because it separates the frequency of wrong episodes from their conditional intensity.

#### 2.4.1. Fluctuations promote frequent transitions to long-duration states where discrimination is better than the deterministic estimate

The significant variability of the ratio *N*_*W*_ */N*_*R*_ during episodes with *EW*^*^ *>* 0 shows that instantaneous specificity displays a rich pattern of fluctuations, as illustrated in Fig. 8b. Thus, the mean value of *ρ*_stoch_ provides only a global measure of discrimination – it does not capture the pattern of favorable and unfavorable intervals that alternate along the stochastic trajectory. To characterize this temporal structure, it is interesting to consider the statistics of excursions where *ρ*(*t*) = *N*_*W*_ (*t*)*/N*_*R*_(*t*) rises above the equilibrium value, *ρ*(*t*) *> ρ*_eq_, or falls below the deterministic prediction, *ρ*(*t*) *< ρ*_det_ *< ρ*_eq_, and the corresponding sojourn times in the two regions. To this end, we define two classes of events, and the relative occurrence times, defined by excursions above/below a given threshold. With upper and lower thresholds *ρ*_eq_ and *ρ*_det_, respectively, this procedure identifies two pairs of sojourn (*τ*) and inter-excursion (Δ*t*) times, as detailed in Table 2. Sojourns below *ρ*_det_ correspond to time stretches in which the readout is transiently more discriminating than predicted by the mean-field description, including states with b*EW*^*^ = 0 at the readout. By contrast, sojourns above *ρ*_eq_ identify highly unfavorable episodes, in which fluctuations transiently increase the wrong-to-correct complex ratio above its equilibrium value. The distributions reported in Fig. 9 show that stochastic excursions to highly favorable discrimination states are frequent and can persist over relatively long timescales, whereas episodes with *ρ*(*t*) *> ρ*_eq_ are substantially shorter and occur less frequently. In particular, the distribution of sojourn times in high-discrimination states (*τ* ^*<*det^) displays a fat tail, indicative of an excess of longer-than-average residence times. The stochastic description contains information that is not accessible from the mean specificity alone. Notably, it appears that fluctuations reshape discrimination through frequent transitions to and long residences in higher-than-deterministic discrimination states.

**Table 2.** Definition of sojourn and inter-excursion times (*k* = 1, 2, …) for events relative to the two specificity thresholds *ρ*_eq_ and *ρ*_det_. Vertical arrows indicate whether the crossing is from below (↑) or from above (↓) with respect to the threshold.

| Event | Sojourn times | Inter-excursion times |
| --- | --- | --- |
| Crossing the upper threshold $\rho_{\text{eq}}$ | $\tau_k^{>\text{eq}} = t_k^{\downarrow\text{eq}} - t_k^{\uparrow\text{eq}}$ | $\Delta t_k^{>\text{eq}} = t_{k+1}^{\uparrow\text{eq}} - t_k^{\uparrow\text{eq}}$ |
| Crossing the lower threshold $\rho_{\text{det}}$ | $\tau_k^{<\text{det}} = t_k^{\uparrow\text{det}} - t_k^{\downarrow\text{det}}$ | $\Delta t_k^{<\text{det}} = t_{k+1}^{\downarrow\text{det}} - t_k^{\downarrow\text{det}}$ |

**Figure 9.**
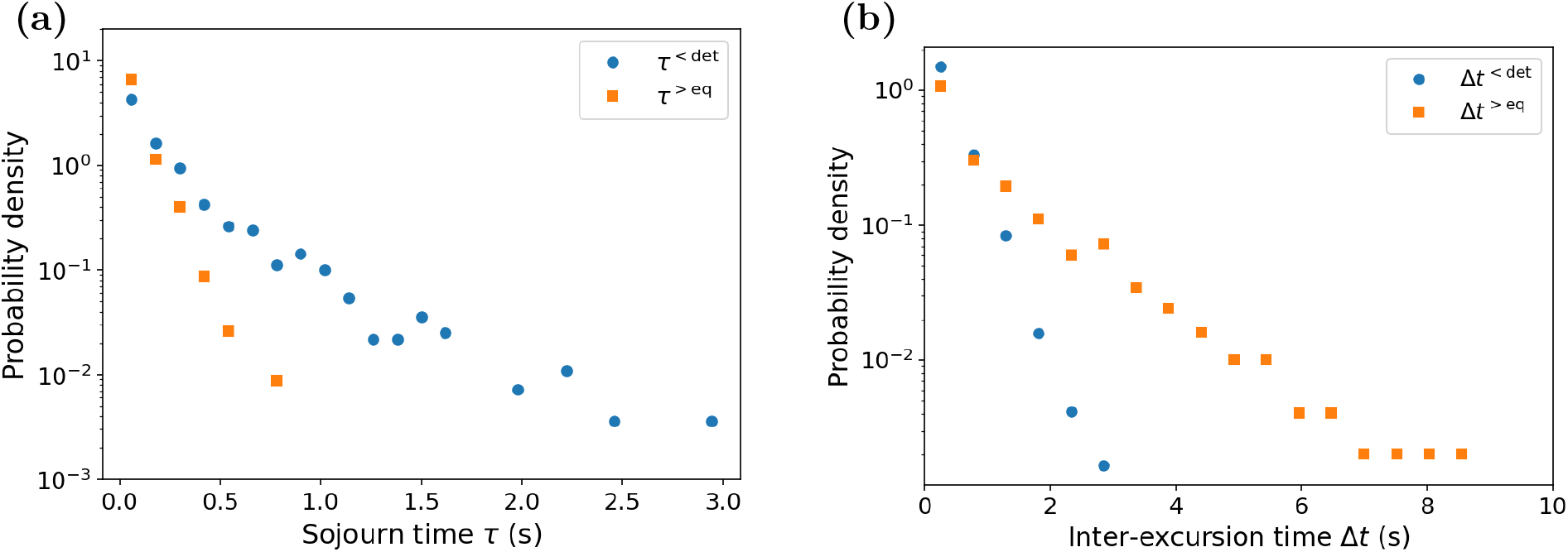
Statistics of stochastic excursions of the instantaneous specificity after the initial transient *t*_0_ = 10^2^ s. **(a)** Probability density of sojourn times *τ*. **(b)** Probability density of inter-excursion times Δ*t* (see also Table 2). Other parameters as in Fig. 2.

## 3. Discussion

In this work we introduced a thermodynamically consistent reaction–diffusion model of spatial proofreading. The main idea is that diffusion provides the necessary delay between successive testing stages where wrong or right substrate can bind to or dissociate from a given enzyme. Of course, transitions associated with such delay should be hard-driven for the system to outperform the accuracy afforded by affinity differences in thermodynamic equilibrium. This is achieved through steady gradients of chemical species. At variance with the original formulation [11], in our model gradients appear self-consistently as a consequence of driving. More specifically, this is realized explicitly through a kinase/phosphatase switch that burns ATP to activate (i.e. phosphorylate) substrate molecules.

Our thermodynamically consistent framework allowed us to determine how the error rate (specificity) depends on thermodynamic driving, gradient confinement, diffusive transport and selective complex dissociation. Our results show that spatial proofreading is effective only within an appropriate balance of different physical timescales: phosphatase activity removes inorganic phosphate from active substrates and hence confines their spatial gradients. Diffusion sets the transport time between the activation and the readout regions, while dissociation rates determine whether the wrong complex is selectively removed before reaching the readout boundary.

We uncover a distinct non-monotonic dependence of specificity on diffusion, with optimal discrimination (minimum specificity) appearing in a well-defined range of diffusion times. This is because diffusion does not simply control the speed of transport, it also defines the temporal window available for non-futile selective dissociation. Hence, optimal discrimination arises as a subtle compromise between diverging forces – transport must be slow enough to allow removal of *EW*^*^ molecules, but not so slow that the readout region becomes effectively isolated from the activation source at the opposite end. Similarly, changing the absolute scale of 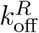and 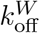 shows that the ratio 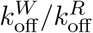 alone is not sufficient to determine discrimination; the relevant condition is the matching between dissociation times and transport times.

An additional constraint on the effectiveness of proofreading is the magnitude of the correct readout. Strong discrimination would be of limited relevance if it were achieved only at the cost of an extremely small concentration of the correct complex, as recently emphasized in the context of activity–fidelity trade-offs [21]. We therefore examined explicitly the relation between specificity and the correct-complex concentration at the readout, [*ER*^*^]_*n*_ (see Supplementary Materials).

Although a compromise between these two quantities emerges in some regimes, particularly for very slow transport, enhanced discrimination is not systematically associated with a vanishing correct readout. In particular, the reference parameter set and many of the sampled parameter sets display *ρ/ρ*_eq_ *<* 1 while maintaining [*ER*^*^]_*n*_ above the 1 nM reference benchmark.

Analyses performed in different 2D geometries further indicate that specificity can be modulated by the geometry of the domain and by the spatial distribution of sources (i.e. kinases). Localized sources generate spatially heterogeneous readout profiles, with weaker discrimination along direct diffusive paths and improved discrimination in regions connected to sources through longer trajectories. This finding might have interesting biological implications: cells could tune molecular accuracy not only by changing reaction rates or enzyme abundances, but also by reorganizing the spatial distribution of kinases, phosphatases and readout regions. In this perspective, geometry and subcellular localization act as additional control parameters for biochemical specificity.

Cells often have to deal with extremely low copy numbers of specific chemical species. The question therefore arises as to the role of intrinsic fluctuations in spatial proofreading mechanisms. Stochastic simulations of our model showed that, in regimes of low molecular occupancy, specificity is not fully captured by average concentrations. A more meaningful measure of accuracy is an event-weighted specificity, which factors in separately (i) the probability of observing a wrong-readout event and (ii) the conditional amplitude of the error when such an event occurs. This distinction is particularly useful in regimes dominated by rare events, where a low average error does not necessarily imply that the instantaneous ratio *N*_*W*_ */N*_*R*_ is small, but rather that wrong-readout episodes are infrequent and short-lived. Furthermore, a more in-depth analysis of specificity fluctuations shows that the system features both excursions in states where discrimination is worse than that afforded by equilibrium (selective affinity), and excursions to discrimination levels better than the corresponding non-equilibrium mean-field estimate. Interestingly, however, the latter excursions are more frequent and give access to comparatively longer-lived states of fluctuation-enhanced discrimination.

Several natural extensions follow from our theoretical framework. First, a fully three-dimensional geometry would provide a more realistic description of cellular organization, especially in systems where membranes, organelles or nuclear regions constrain diffusion and localization. Second, it would be important to study how spatial proofreading interacts with other nonequilibrium discrimination mechanisms, such as classical kinetic proofreading [3] or catalytic discrimination [8]. In real biochemical networks, these mechanisms are unlikely to act in isolation, and their combined effect could generate regimes of enhanced accuracy or new trade-offs between specificity, speed and dissipation. Finally, future work should connect the model to concrete biological systems. Signaling pathways based on phosphorylation cascades, such as MAPK networks [22], provide a particularly relevant context because they combine kinase–phosphatase cycles, spatial localization, scaffolding and compartmentalized signal propagation.

Overall, our results show that spatial proofreading is controlled by the coupling between nonequilibrium driving, diffusive transport and spatial organization. Specificity is therefore not set only by local binding free energies or chemical rate constants, but also by the time and geometry of transport between activation and readout regions. Spatial proofreading may therefore provide a general mechanism by which cells exploit diffusion and spatial organization to enhance molecular discrimination. In order to delve deeply into the design principles of space-embedded molecular error management, these phenomena should be investigated through carefully designed thermodynamically consistent models.

## 4. Materials and Methods

All deterministic and stochastic simulations were performed using the Python package STReNGTHS [23], a computational framework designed to simulate complex reaction–diffusion systems. The reaction network, spatial grid, chemostats and kinase localization patterns were implemented within the same discretized reaction–diffusion framework used throughout the Results.

The deterministic dynamics was obtained by integrating the reaction–diffusion equations associated with the local reaction network described in Tab. 1 for 10^3^ s. We have verified that this is enough to reach convergence to the steady state in all cases considered. Stochastic simulations were performed using the Gillespie algorithm implemented in STReNGTHS on the same discretized reaction–diffusion network. Molecular copy numbers were sampled in the readout compartment, and an initial transient *t*_0_ = 10^2^ s was excluded from all stochastic analyses.

For the standard stochastic regime, trajectories were simulated up to *T*_sim_ = 10^3^ s and sampled using *N*_samp_ = 2 *×* 10^4^ points. In the rare-event regime, where the presence of *EW*^*^ in the readout is strongly suppressed, the simulation time was increased to *T*_sim_ = 10^5^ s and the number of sampled points to *N*_samp_ = 2 *×* 10^6^, in order to collect a sufficient number of wrong-readout episodes. In both cases, this corresponds to a sampling interval 5 *×* 10^*−*2^ s.

## Author contributions

TR, TF, and FP conceived the model. TR performed the simulations and analyzed the results. All authors discussed the results and contributed to writing the manuscript.

## Data availability

All data are available from the authors upon reasonable request.

## Supplementary materials

Supplementary electronic material available online.

## Acknowledgements

The authors would like to thank Vahe Galstyan, Arvind Murugan, Rob Phillips and Alessandro Barducci for a critical reading of this paper and many insightful comments.

1 In practice, we have adjusted *v*_*k*_ so that the overall stationary flux of activated substrate stayed the same irrespective of the different number of sources (i.e. bands with active kinases)

2 This value corresponds to the angular mean of *ρ*(*θ*)*/ρ*_eq_ computed for the uniform peri-nuclear kinase distribution. This averaging is introduced because the reference profile is not perfectly flat: small residual oscillations arise from the discrete geometry of the numerical domain.

## Notes

### Competing Interest Statement

The authors have declared no competing interest.

### Summary of Updates

Expanded text. New section in the supplementary material document.

