## Supplementary material for "A Thermodynamically Consistent Reaction–Diffusion Model of Spatial Proofreading in One and Two Dimensions"

### Supplementary Materials for "A Thermodynamically Consistent Reaction–Diffusion Model of Spatial Proofreading in One and Two Dimensions"

Tommaso Rossi, Thibault Fillion and Francesco Piazza

**6 Activity–specificity trade-off** **21**

#### 1. Thermodynamic constraints and parametrization of energetic chemostats

##### 1.1. Detailed balance at equilibrium

To ensure the thermodynamic consistency of the phosphorylation–dephosphorylation cycle, we impose that the ratio between the product of the rates along one direction of the cycle and the product of the rates along the opposite direction is fixed by the chemical affinity of the cycle. Introducing $\Pi^+$ and $\Pi^-$ as the products of the transition rates along the cycle, respectively in the positive direction and in the opposite direction, the thermodynamic constraint can be written as

$$\frac{\Pi^+}{\Pi^-} = e^{\beta\Delta\mu}, \qquad \beta = \frac{1}{k_B T}. \tag{S1}$$

At thermodynamic equilibrium, namely for $\Delta\mu = 0$, this ratio becomes equal to one and the cycle satisfies detailed balance, according to Kolmogorov's criterion.

In our model, we choose as the positive direction of the cycle the sequence of reactions in which the substrate is first phosphorylated and then dephosphorylated:

$$S + \mathrm{ATP} \to S^* + \mathrm{ADP}, \qquad S^* \to S + P_\mathrm{i}.$$

The reactions followed in the opposite direction are instead

$$S^* + \mathrm{ADP} \to S + \mathrm{ATP}, \qquad S + P_\mathrm{i} \to S^*.$$

We denote by $k_k^+$ and $k_k^-$ the microscopic constants associated respectively with phosphorylation and with the reverse reaction catalyzed by the kinase. Similarly, we denote by $k_p^-$ the constant associated with dephosphorylation and by $k_p^+$ the one associated with the reverse reaction catalyzed by the phosphatase. The ratio between the products of the rates can therefore be written as

$$\frac{\Pi^+}{\Pi^-} = \frac{k_k^+[\mathrm{ATP}]}{k_k^-[\mathrm{ADP}]}\frac{k_p^-}{k_p^+[P_\mathrm{i}]}.$$

The thermodynamic consistency condition therefore becomes

$$\frac{k_k^+[\mathrm{ATP}]}{k_k^-[\mathrm{ADP}]}\frac{k_p^-}{k_p^+[P_\mathrm{i}]} = e^{\beta\Delta\mu}.$$

At thermodynamic equilibrium ($\Delta\mu = 0$), this condition reduces to

$$\frac{k_k^+}{k_k^-}\frac{k_p^-}{k_p^+} = \frac{[\mathrm{ADP}]_\mathrm{eq}[P_\mathrm{i}]_\mathrm{eq}}{[\mathrm{ATP}]_\mathrm{eq}}.$$

We now define the microscopic equilibrium constants of the two catalytic steps and the association equilibrium constant of the chemostats as

$$K_k := \frac{k_k^+}{k_k^-}, \qquad K_p := \frac{k_p^+}{k_p^-}, \qquad K_\mathrm{ATP} := \frac{[\mathrm{ATP}]_\mathrm{eq}}{[\mathrm{ADP}]_\mathrm{eq}[P_\mathrm{i}]_\mathrm{eq}}. \tag{S2}$$

Substituting these definitions into the equilibrium condition gives

$$\frac{K_k}{K_p} = \frac{1}{K_\mathrm{ATP}}, \qquad K_k = \frac{K_p}{K_\mathrm{ATP}}.$$

This constraint shows that $K_k$ and $K_p$ are not independent, but that only their ratio is fixed by the detailed-balance condition at equilibrium. For simplicity, we set $K_k = 1$, which gives $K_p = K_\mathrm{ATP}$. This parametrization ensures that the cycle satisfies detailed balance at equilibrium, while out of equilibrium the driving is controlled by the chemostat concentrations through the following relation:

$$\Delta\mu = k_B T \ln\left(\frac{[\mathrm{ATP}]\,[\mathrm{ADP}]_\mathrm{eq}\,[P_\mathrm{i}]_\mathrm{eq}}{[\mathrm{ATP}]_\mathrm{eq}\,[\mathrm{ADP}]\,[P_\mathrm{i}]}\right). \tag{S3}$$

##### 1.2. Physiological reference values

The physiological concentrations of ATP, ADP and $P_\mathrm{i}$ are taken as representative of a generic human cell. The concentrations of ATP and ADP are computed as the average of the concentrations measured in human cells (erythrocytes and lymphocytes for ATP; eosinophils, erythrocytes, lymphocytes and monocytes for ADP) reported in [1] (Tab. 8).

$$\begin{aligned}[\mathrm{ATP}]_c &= \langle\{1519,\ 1629,\ 1871,\ 390,\ 463,\ 1900,\ 2303\}\rangle \quad \mu\mathrm{M} = \\ &= 1439.3\,\mu\mathrm{M}.\end{aligned} \tag{S4}$$

$$\begin{aligned}[\mathrm{ADP}]_c &= \langle\{22,\ 171,\ 191,\ 871,\ 230,\ 16,\ 212,\ 360,\ 32,\ 24\}\rangle \quad \mu\mathrm{M} = \\ &= 212.9\,\mu\mathrm{M}.\end{aligned} \tag{S5}$$

The physiological concentration of inorganic phosphate is taken from [1] (Tab. 6, human lymphocytes):

$$[P_\mathrm{i}]_c = 2770 \quad \mu\mathrm{M}. \tag{S6}$$

##### 1.3. Chemical equilibrium and determination of $K_\mathrm{ATP}$

From [2], we take the standard Gibbs free energy for the hydrolysis of $ATP$ into ADP and $P_\mathrm{i}$ to be $\Delta G_h^\circ = -36.53$ kJ/mol (Tab. 4.1 at $T = 298.15$ K, ionic strength 0.10 M and pH 7). We define the association equilibrium constant for the reaction ADP + $P_\mathrm{i} \rightleftharpoons$ ATP as

$$K_\mathrm{ATP} = \exp\left(\frac{\Delta G_h^\circ}{RT}\right) = 3.9831 \times 10^{-7}\,\mathrm{M}^{-1} \tag{S7}$$

where $R = 8.314\ \mathrm{J\,mol^{-1}\,K^{-1}}$ is the ideal gas constant.

The equilibrium concentrations of ATP, ADP and $P_\mathrm{i}$, corresponding to the physiological quantities defined in the previous section, can be obtained as follows: we introduce the total concentrations

$$\begin{aligned}[\mathrm{ADP}]_\mathrm{tot} &= [\mathrm{ADP}]_c + [\mathrm{ATP}]_c, \\ [P_\mathrm{i}]_\mathrm{tot} &= [P_\mathrm{i}]_c + [\mathrm{ATP}]_c,\end{aligned} \tag{S8}$$

and assume that these quantities are conserved, namely constant in time.

From the definition of $K_\mathrm{ATP}$, we obtain

$$K_\mathrm{ATP} = \frac{[\mathrm{ATP}]_\mathrm{eq}}{[\mathrm{ADP}]_\mathrm{eq}[P_\mathrm{i}]_\mathrm{eq}} = \frac{[\mathrm{ATP}]_\mathrm{eq}}{\big([\mathrm{ADP}]_\mathrm{tot} - [\mathrm{ATP}]_\mathrm{eq}\big)\big([P_\mathrm{i}]_\mathrm{tot} - [\mathrm{ATP}]_\mathrm{eq}\big)}. \tag{S9}$$

This relation can be rewritten as a second-degree polynomial in $[\mathrm{ATP}]_\mathrm{eq}$:

$$-K_\mathrm{ATP}[\mathrm{ATP}]_\mathrm{eq}^2 + [\mathrm{ATP}]_\mathrm{eq}\,(1 + K_\mathrm{ATP}[\mathrm{ADP}]_\mathrm{tot} + K_\mathrm{ATP}[P_\mathrm{i}]_\mathrm{tot}) - K_\mathrm{ATP}[\mathrm{ADP}]_\mathrm{tot}[P_\mathrm{i}]_\mathrm{tot} = 0. \tag{S10}$$

The physically relevant root, which satisfies the condition $0 < [\mathrm{ATP}]_\mathrm{eq} < \min\left([\mathrm{ADP}]_\mathrm{tot}, [P_\mathrm{i}]_\mathrm{tot}\right)$, is given by

$$[ATP]_\mathrm{eq} = 2.7698 \times 10^{-6}\,\mu\mathrm{M}. \tag{S11}$$

The equilibrium concentrations of ADP and $P_\mathrm{i}$ are obtained by subtracting the equilibrium value of ATP from the corresponding total concentration:

$$[\mathrm{ADP}]_\mathrm{eq} = [\mathrm{ADP}]_\mathrm{tot} - [\mathrm{ATP}]_\mathrm{eq}, \qquad [P_\mathrm{i}]_\mathrm{eq} = [P_\mathrm{i}]_\mathrm{tot} - [\mathrm{ATP}]_\mathrm{eq}. \tag{S12}$$

##### 1.4. Chemostats as a function of the driving $\beta\Delta\mu$

In order to obtain the chemostat concentrations as a function of $\beta\Delta\mu$, we introduce $Q := \exp(\beta\Delta\mu)$, which quantifies the strength of the driving. The thermodynamic constraint can be written as

$$\frac{[\mathrm{ATP}]}{[\mathrm{ATP}]_{\mathrm{eq}}} \frac{[\mathrm{ADP}]_{\mathrm{eq}}}{[\mathrm{ADP}]} \frac{[P_{\mathrm{i}}]_{\mathrm{eq}}}{[P_{\mathrm{i}}]} = Q. \tag{S13}$$

Assuming that $[\mathrm{ADP}]_{\mathrm{tot}}$ and $[P_{\mathrm{i}}]_{\mathrm{tot}}$ are conserved, we have

$$[\mathrm{ADP}] = [\mathrm{ADP}]_{\mathrm{tot}} - [\mathrm{ATP}], \qquad [P_{\mathrm{i}}] = [P_{\mathrm{i}}]_{\mathrm{tot}} - [\mathrm{ATP}]. \tag{S14}$$

Substituting Eq. (S14) into Eq. (S13) and solving for $[ATP]$ (analogously to Eq. (S9)) gives

$$A := \frac{1}{K_{\mathrm{ATP}}} + Q([\mathrm{ADP}]_{\mathrm{tot}} + [P_{\mathrm{i}}]_{\mathrm{tot}})\,, \tag{S15}$$

$$\begin{aligned} [\mathrm{ATP}] &= \frac{A - \sqrt{A^2 - 4Q^2[\mathrm{ADP}]_{\mathrm{tot}}[P_{\mathrm{i}}]_{\mathrm{tot}}}}{2Q} \\ &= \frac{2Q[\mathrm{ADP}]_{\mathrm{tot}}[P_{\mathrm{i}}]_{\mathrm{tot}}}{A + \sqrt{A^2 - 4Q^2[\mathrm{ADP}]_{\mathrm{tot}}[P_{\mathrm{i}}]_{\mathrm{tot}}}} \end{aligned} \tag{S16}$$

where the physically relevant root has been chosen. The concentrations of [ADP] and $[P_{\mathrm{i}}]$ are obtained from Eq. (S14).

Figure S1 shows the behavior of the concentrations of the three chemostats as a function of $\beta\Delta\mu$. For $\beta\Delta\mu \gtrsim 1$, [ATP] increases monotonically from very small values, of the order of $10^{-5}$ $\mu$M, until it reaches a plateau for $\beta\Delta\mu \gtrsim 20$ of approximately $10^3$ $\mu$M. Conversely, [ADP] and $[P_{\mathrm{i}}]$ decrease as the driving increases: [ADP] decreases by many orders of magnitude down to values of the order of $10^{-5}$ $\mu$M, while $[P_{\mathrm{i}}]$ changes more moderately, remaining of the same order of magnitude. Moreover, the correct recovery of the equilibrium values in the case $\beta\Delta\mu = 0$ provides a check of the consistency of the adopted parametrization procedure.

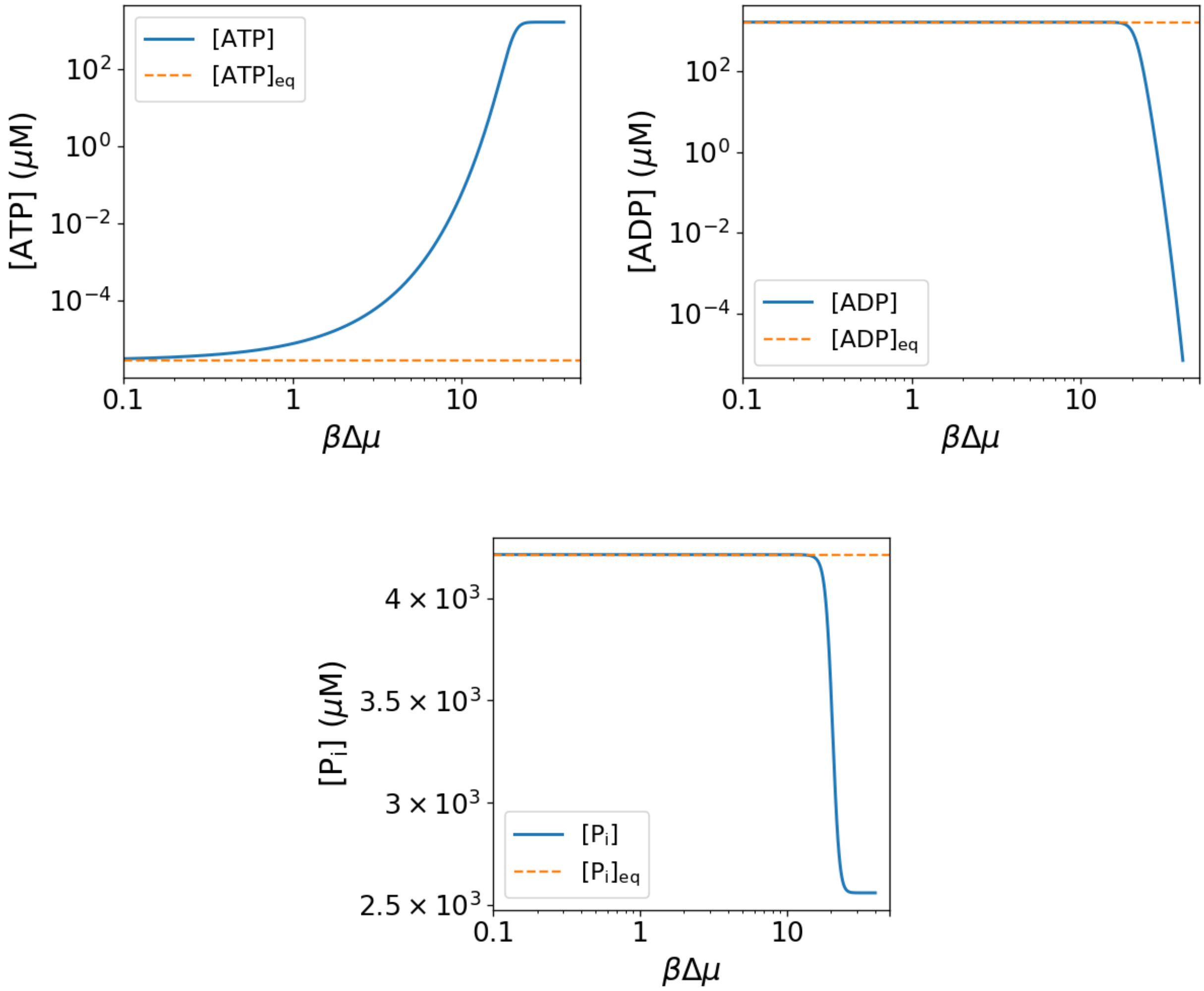

**Figure S1:** Concentrations of the chemostats as a function of $\beta\Delta\mu \in [0.1, 40]$.

#### 2. Reference setup and parameter choice

Here we report the reference setup used in the simulations. The energetic chemostats and thermodynamic constants are summarized in Tab. S1; the initial concentrations are reported in Tab. S2; and the kinetic, geometric and transport parameters are listed in Tab. S3.

**Table S1:** Energetic chemostats and thermodynamic constants.

| Parameter | Value | Unit | Description |
|---|---|---|---|
| $[ATP]_c$ | $1.4393 \times 10^3$ | $\mu$M | cytosolic ATP concentration |
| $[ADP]_c$ | $2.129 \times 10^2$ | $\mu$M | cytosolic ADP concentration |
| $[P_i]_c$ | $2.770 \times 10^3$ | $\mu$M | cytosolic $P_\mathrm{i}$ concentration |
| $K_{\mathrm{ATP}}$ | $3.9831 \times 10^{-13}$ | $\mu\mathrm{M}^{-1}$ | equilibrium constant |
| $[ATP]_{\mathrm{eq}}$ | $2.7698 \times 10^{-6}$ | $\mu$M | ATP concentration at equilibrium |

The choice of the physiological values reported in Tab. S1, the determination of the equilibrium concentrations, and the procedure used to impose the energetic chemostats as functions of the driving $\beta\Delta\mu$ are discussed in detail in Sec. 1.

**Table S2:** Initial concentrations.

| Parameter | Value | Unit | Description |
|---|---|---|---|
| $[W_0]$ | 3.0 | $\mu$M | initial concentration of $W$ |
| $[R_0]$ | 3.0 | $\mu$M | initial concentration of $R$ |
| $[E_0]$ | 0.60 | $\mu$M | initial concentration of $E$ |

The initial concentrations of the two substrates are assumed to be equal, $[W_0] = [R_0] = 3.0\,\mu$M, while the initial enzyme concentration is fixed to $[E_0] = 0.60\,\mu$M, so as to consider a biologically plausible condition in which the enzyme is less abundant than the substrates [3]. In particular, the simulations are initialized with an enzyme-to-substrate ratio of approximately $1 : 5$ for each substrate. This choice avoids the enzyme-excess limit and allows enzyme sequestration by the complexes to affect the free-enzyme profile. Moreover, these values lie within typical cellular orders of magnitude and allow us to study the spatial proofreading mechanism both in the deterministic regime and in the stochastic regime.

**Table S3:** Reference kinetic, geometric and transport parameters.

| Parameter | Value | Unit | Description |
|---|---|---|---|
| $n$ | 10 | – | number of compartments |
| $\Delta x$ | 1 | $\mu$m | spatial discretization step |
| $L$ | 10 | $\mu$m | source–readout distance |
| $D$ | 10 | $\mu\mathrm{m}^2\,\mathrm{s}^{-1}$ | diffusion coefficient |
| $k_{\mathrm{on}}$ | $10^6$ | $\mathrm{M}^{-1}\mathrm{s}^{-1}$ | enzyme–substrate association rate |
| $k_{\mathrm{off}}^R$ | 0.1 | $\mathrm{s}^{-1}$ | correct-complex dissociation rate |
| $k_{\mathrm{off}}^W$ | 1 | $\mathrm{s}^{-1}$ | wrong-complex dissociation rate |
| $v_k$ | $3.0 \times 10^2$ | $\mathrm{M}^{-1}\mathrm{s}^{-1}$ | kinase kinetic scale |
| $v_p$ | 5 | $\mathrm{s}^{-1}$ | phosphatase kinetic scale |
| $\beta\Delta\mu$ | 20 | – | nonequilibrium driving |

Considering $n = 10$ and $\Delta x = 1\,\mu$m, the total length of the one-dimensional domain is $L = n\Delta x = 10\,\mu$m, in agreement with the original spatial proofreading model [4]. The association and dissociation rates, $k_{\mathrm{on}}$, $k_{\mathrm{off}}^R$ and $k_{\mathrm{off}}^W$, are set equal to the reference values used in the original spatial proofreading framework [4]. For equal association rates and equal initial substrate concentrations, this gives the equilibrium specificity $\rho_{\text{eq}} = k_{\text{off}}^R / k_{\text{off}}^W = 0.1$.

The kinetic parameters $v_k$ and $v_p$ are chosen so that the effective kinase and phosphatase rates are comparable with those used in the original model. In particular, the reference values correspond to effective rates of the order $k_k \simeq 0.2\,\text{s}^{-1}$ and $k_p \simeq 5\,\text{s}^{-1}$. Operationally, these rates can be interpreted as net reaction fluxes normalized by the concentration of the corresponding reactant:

$$k_k = \frac{v_k[\text{ATP}][S] - v_k[\text{ADP}][S^*]}{[S]}, \tag{S17}$$

$$k_p = \frac{v_p[S^*] - v_p K_{\text{ATP}}[P_i][S]}{[S^*]}. \tag{S18}$$

Here, the substrate concentrations are evaluated at steady state.

Among the parameters directly comparable with the original spatial proofreading model, the main reference value chosen differently is the diffusion coefficient $D$. In Ref. [4], the reference value was $D = 1\,\mu\text{m}^2\,\text{s}^{-1}$, whereas here we use $D = 10\,\mu\text{m}^2\,\text{s}^{-1}$, which remains a biologically plausible value for protein-sized species in the cytoplasm [3]. The reason for this choice is that diffusion directly controls the transport time between the activation and readout regions and therefore the time window available for selective dissociation. In particular, slow diffusion can strongly reduce $\rho$ and make the spatial proofreading mechanism very efficient. We therefore adopt a reference value of $D$ that is not too small, so that the role of the other kinetic parameters can also be clearly resolved.

Finally, the reference value of the driving is set to $\beta\Delta\mu = 20$, which is of the order expected under cellular nucleotide conditions [3]. As shown above, this value corresponds to a strongly nonequilibrium regime in which the phosphorylation–dephosphorylation cycle can sustain a stable active-substrate gradient.

#### 3. Additional analysis of nonequilibrium driving

##### 3.1. Reference driving sweep

In this paragraph, we analyze in more detail the response of the model to the nonequilibrium driving $\beta\Delta\mu$ in the reference one-dimensional parameter set. Fig. S2 shows both the resulting normalized specificity and the corresponding steady-state concentrations of the complexes in the readout compartment.

For weak driving, $\beta\Delta\mu \lesssim 3$, the system remains close to equilibrium: the phosphorylation–dephosphorylation cycle does not sustain a pronounced active-substrate gradient and the concentrations of the complexes in the readout compartment remain small. As a consequence, the spatial separation between activation and readout cannot be efficiently exploited and $\rho/\rho_{eq} \simeq 1$.

As the driving is increased, the concentration of activated substrates reaching the readout region grows. In the intermediate range, approximately $\beta\Delta\mu \simeq$ 7–10, the nonequilibrium cycle becomes sufficiently strong to maintain a spatial gradient of active substrates. In this regime, the transport delay becomes effective because wrong complexes are more likely to dissociate before reaching the readout compartment, leading to a substantial decrease in the normalized specificity.

For larger values of the driving, $\beta\Delta\mu \gtrsim 20$, both $[ER^*]_n$ and $[EW^*]_n$ approach a plateau: increasing $\beta\Delta\mu$ further does not significantly change the amount of activated complexes reaching the readout region and $\rho/\rho_{\text{eq}}$ correspondingly approaches an asymptotic value. This indicates that, beyond a sufficiently large driving, thermodynamic forcing is no longer the limiting factor for discrimination. Most of the gain in normalized specificity is already obtained for moderate values of the driving, around $\beta\Delta\mu \simeq$ 10–15, while the high-driving regime is mainly controlled by kinetic and transport parameters.

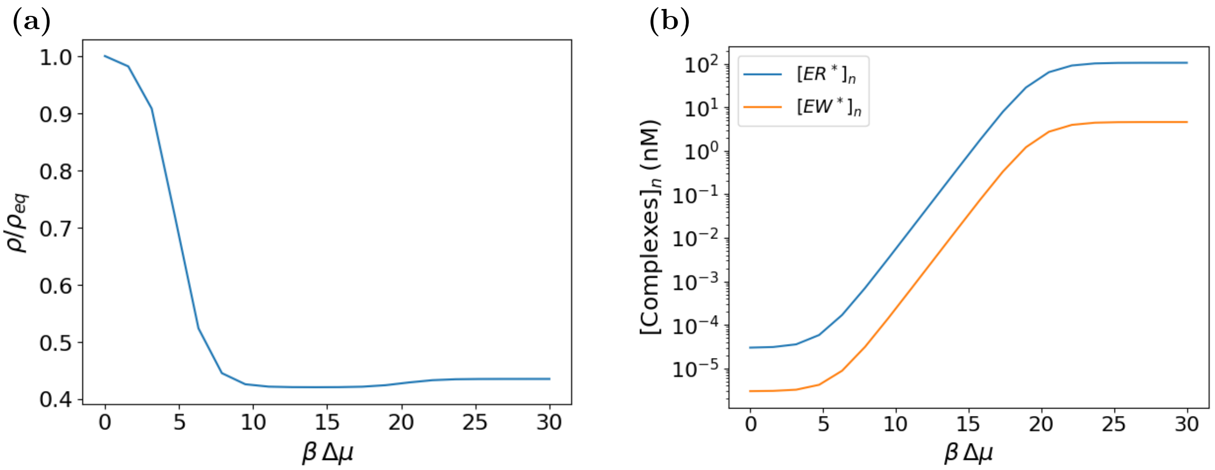

**Figure S2:** Response of the reference one-dimensional system to the nonequilibrium driving $\beta\Delta\mu$. **(a)** Normalized specificity $\rho/\rho_{\mathrm{eq}}$. **(b)** Steady-state concentrations of the complexes $[ER^*]_n$ and $[EW^*]_n$ in the readout compartment. All other parameters are fixed to the reference values reported in Sec. 2.

The slight increase of $\rho/\rho_{\mathrm{eq}}$ observed at high driving, after the minimum reached at intermediate $\beta\Delta\mu$, shows that increasing the thermodynamic force does not always lead to a monotonic improvement in discrimination. In fact, at intermediate driving, the active-substrate gradient is sufficiently maintained and the spatial transport delay can efficiently act as a selective filter, allowing the wrong complex to dissociate more frequently before readout. When the driving is further increased, the system approaches a high-driving plateau, where the amount of activated substrate reaching the readout becomes only weakly sensitive to further changes in the chemostat concentrations and the specificity is mainly controlled by kinetic and transport parameters.

##### 3.2. Spatial profiles of activated substrates at different driving

The steady-state spatial profiles of the activated substrates $R^*$ and $W^*$ are shown in Fig. S3 for different values of $\beta\Delta\mu$. These profiles directly illustrate how the nonequilibrium driving sustains the active-substrate gradient. For small driving, close to equilibrium, the concentrations of activated substrates remain very low and the profiles are nearly flat, as expected when the phosphorylation–dephosphorylation cycle is unable to maintain a pronounced spatial gradient. As $\beta\Delta\mu$ is increased, a gradient is progressively established, with a maximum in the first compartment, where kinase activity is localized, and a monotonic decrease toward the readout compartment.

At intermediate and large driving, increasing $\beta\Delta\mu$ mainly raises the overall concentration of activated substrates, while the profile shape changes only weakly. The profiles therefore shift upward without appreciable changes in their spatial decay, indicating that the system progressively approaches a high-driving regime. No significant qualitative difference is observed between $R^*$ and $W^*$, consistently with the fact that the two substrates are activated and deactivated with identical kinetic rates. As a consequence, the spatial profiles shown here should be interpreted as the common nonequilibrium substrate gradient on which proofreading acts, whereas discrimination between correct and wrong substrates arises primarily from the different dissociation rates of the corresponding complexes, $ER^*$ and $EW^*$.

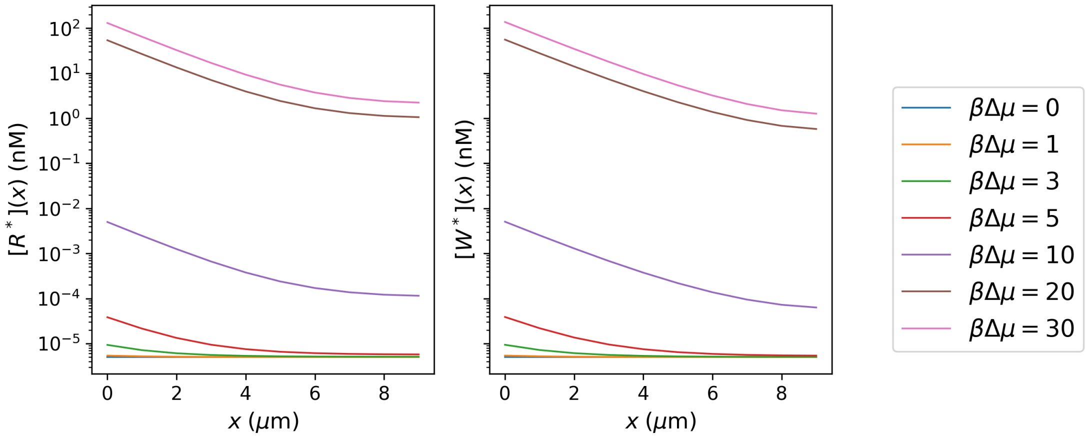

**Figure S3:** Steady-state spatial profiles of the activated substrates, $[R^*](x)$ and $[W^*](x)$, for different values of the nonequilibrium driving $\beta\Delta\mu$. All other parameters are fixed to the reference values reported in Sec. 2.

##### 3.3. Driving response for different kinetic and transport regimes

We then asked whether the response to the nonequilibrium driving is universal or depends on the kinetic and transport regime. To this end, we repeated the sweep in $\beta\Delta\mu$ while varying separately the phosphatase rate $v_p$, the diffusion coefficient $D$ and the absolute scale of the dissociation rates at fixed ratio $\alpha = k_{\text{off}}^W/k_{\text{off}}^R$. The results are shown in Fig. S4.

In all cases, for small values of $\beta\Delta\mu$ the system remains close to the equilibrium limit, whereas increasing the driving initially improves discrimination. However, the high-driving value of $\rho/\rho_{\text{eq}}$ is not universal, but depends strongly on the kinetic and transport parameters: increasing the phosphatase rate $v_p$ lowers the high-driving plateau of $\rho/\rho_{\text{eq}}$, consistently with the role of the phosphatase in confining the active-substrate gradient. At the same time, larger values of $v_p$ shift the onset of the improvement toward higher driving, because stronger dephosphorylation requires a larger thermodynamic force to sustain a sufficient amount of activated substrate at the readout. Reducing $D$ generally makes discrimination more effective, showing that the same thermodynamic driving can produce very different levels of discrimination depending on the diffusive transport time $\tau_D = L^2/D$. Finally, faster dissociation rates lead to a lower high-driving plateau of $\rho/\rho_{\text{eq}}$, whereas slower dissociation rates keep the system closer to the equilibrium limit.

The non-monotonic dependence on the driving is clearly visible in parameter regimes where spatial proofreading remains relatively inefficient, with $\rho/\rho_{\text{eq}}$ close to one. In these cases, intermediate values of $\beta\Delta\mu$ provide a finite gain in discrimination by sustaining an active-substrate gradient, but further increasing the driving cannot compensate for unfavorable transport or dissociation times. The system therefore approaches a high-driving plateau in which the specificity is set mainly by kinetic and spatial parameters rather than by thermodynamic forcing. By contrast, when proofreading is already highly efficient and $\rho/\rho_{\text{eq}}$ is strongly reduced, the wrong complex is depleted so effectively during transport that the high-driving increase becomes negligible. In this regime, increasing the driving mainly maintains the active-substrate supply, while the ratio between wrong and correct complexes is controlled by selective dissociation. As a result, the curve decreases and then saturates at a low value, without displaying an appreciable minimum followed by a rise.

Overall, these additional sweeps show that a sufficiently large nonequilibrium driving is necessary to maintain the active-substrate gradient, but it is not sufficient to determine the final specificity. Once the system is far from equilibrium, the level of discrimination is set by the interplay between gradient confinement, diffusive transport and complex dissociation.

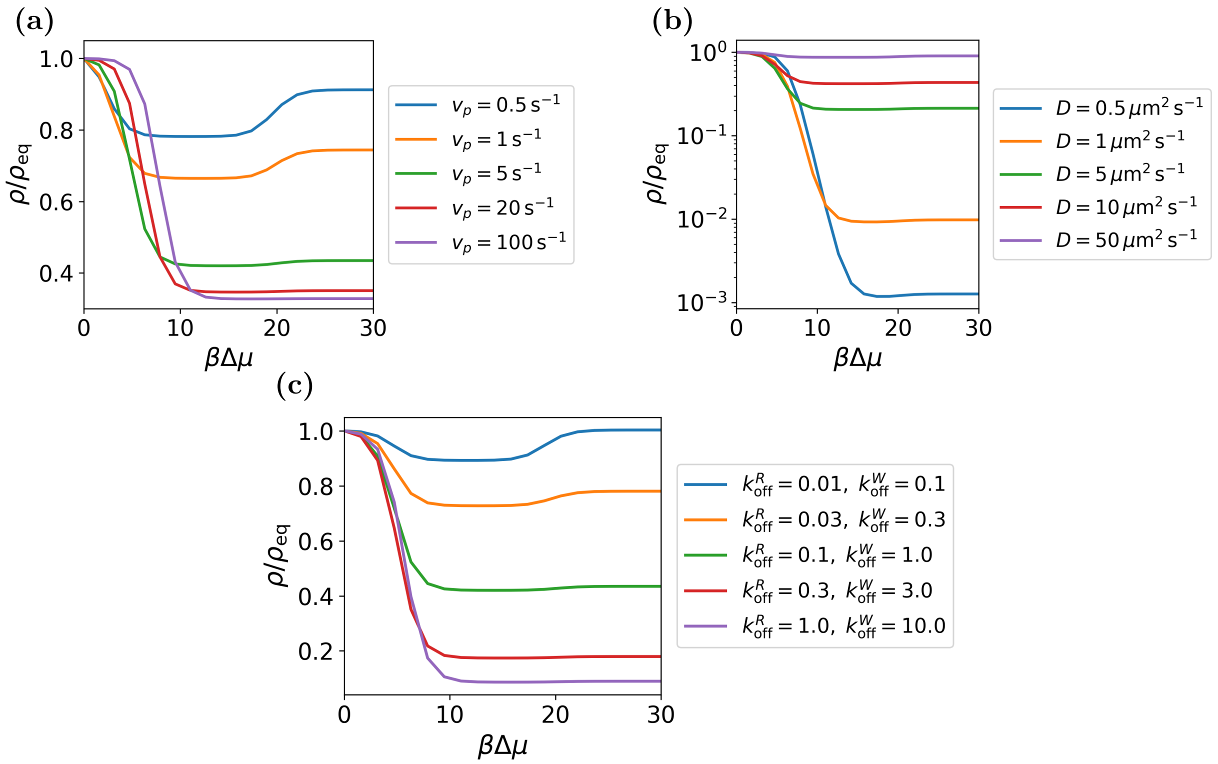

**Figure S4:** Response of the normalized specificity $\rho/\rho_{\text{eq}}$ to the nonequilibrium driving $\beta\Delta\mu$ in different kinetic and transport regimes. **(a)** Driving response for different values of the phosphatase rate $v_p$. **(b)** Driving response for different values of the diffusion coefficient $D$. **(c)** Driving response for different absolute scales of the dissociation rates, at fixed ratio $k_{\text{off}}^W/k_{\text{off}}^R = 10$. The remaining parameters are fixed to the reference values reported in Sec. 2.

#### 4. Additional one-dimensional parameter sweeps

##### 4.1. Effect of rebinding: $k_{\text{on}}$ and $[E_0]$

In the original work on spatial proofreading, the role of rebinding was analyzed through the effective parameter

$$k_b = k_{\text{on}}\rho_E, \tag{S19}$$

where $\rho_E$ denotes the free enzyme density, assumed to be spatially uniform in the minimal model of Ref. [4]. This parameter sets the effective rate at which a substrate can rebind the enzyme after dissociation. Increasing $k_b$ reduces the time spent by the substrate in the free state and can therefore suppress the proofreading advantage. In our model, $k_{\text{on}}$ and the total enzyme concentration $[E_0]$ are independent parameters. This allows us to disentangle the contribution of the microscopic association rate from that of the total enzyme abundance.

First, we varied $k_{\text{on}}$ in the range $10^5$–$10^{10}\,\text{M}^{-1}\text{s}^{-1}$, while keeping the other parameters fixed. As shown in Fig. S5a, for small values of $k_{\text{on}}$ the normalized specificity remains approximately constant, with $\rho/\rho_{\text{eq}} \simeq 0.4$. In this regime, enzyme–substrate association is not sufficiently fast to alter the proofreading dynamics significantly. As $k_{\text{on}}$ increases, rebinding becomes progressively more efficient: after dissociation, activated substrates are rapidly recaptured before being deactivated by the phosphatase. This reduces the time window in which dissociation can selectively penalize the wrong substrate and leads to an increase of $\rho/\rho_{\text{eq}}$, which can eventually reach values larger than one.

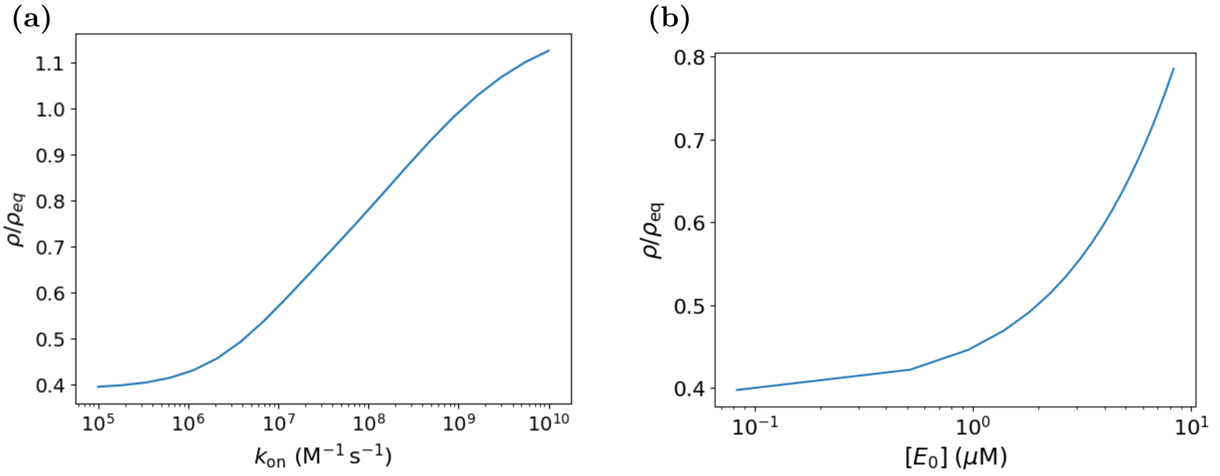

**Figure S5:** **(a)** Normalized specificity $\rho/\rho_{\mathrm{eq}}$ as a function of the association rate $k_{\mathrm{on}}$. **(b)** Normalized specificity $\rho/\rho_{\mathrm{eq}}$ as a function of the total enzyme concentration $[E_0]$. The remaining parameters are fixed to the reference values reported in Sec. 2.

We then varied the total enzyme concentration $[E_0]$ in the range $8.30\times10^{-2}$–$8.30\,\mu$M. As shown in Fig. S5b, increasing $[E_0]$ produces a monotonic increase of $\rho/\rho_{\mathrm{eq}}$ over the range considered. This behavior can again be interpreted in terms of rebinding: a larger enzyme abundance increases the probability that a dissociated activated substrate rapidly forms a new complex. As a result, the wrong substrate $W^*$, instead of remaining free long enough to be deactivated by the phosphatase, can undergo repeated binding events and contribute more efficiently to the formation of $EW^*$ in the readout compartment.

Overall, both sweeps confirm the same physical mechanism, namely that increasing either $k_{\mathrm{on}}$ or $[E_0]$ decreases the characteristic rebinding time,

$$t_{\mathrm{on}} \sim \frac{1}{k_{\mathrm{on}}[E]}, \tag{S20}$$

thereby reducing the time interval during which the wrong substrate can be selectively rejected after dissociation. These results are consistent with the role of rebinding discussed in Ref. [4], while extending the analysis to a model in which $k_{\mathrm{on}}$, $[E_0]$ and the spatial distribution of free enzyme are treated explicitly.

##### 4.2. Effect of kinase activation rate $v_k$

We next analyzed the effect of the kinase activation rate $v_k$, which controls the production of activated substrates in the first compartment. To this end, we performed a logarithmic sweep of $v_k$ over the range 0.60–6.02 $\times\, 10^5\,\mathrm{M}^{-1}\mathrm{s}^{-1}$. The result is shown in Fig. S6a.

In contrast to the phosphatase rate $v_p$, the effect of $v_k$ on the normalized specificity is weak over the range explored. Although $v_k$ is varied over five orders of magnitude, $\rho/\rho_{\mathrm{eq}}$ changes only from approximately 0.42 to 0.48. This weak dependence is consistent with the role of the kinase in the model: activation acts symmetrically on correct and wrong substrates and therefore does not introduce a selective mechanism comparable to the differential dissociation of the complexes.

Changing $v_k$ mainly modifies the overall production flux of activated substrates entering the domain. As shown in Fig. S6b, increasing $v_k$ increases the concentrations of both complexes in the readout compartment, but it does not substantially alter their ratio. This explains why the specificity is only weakly affected.

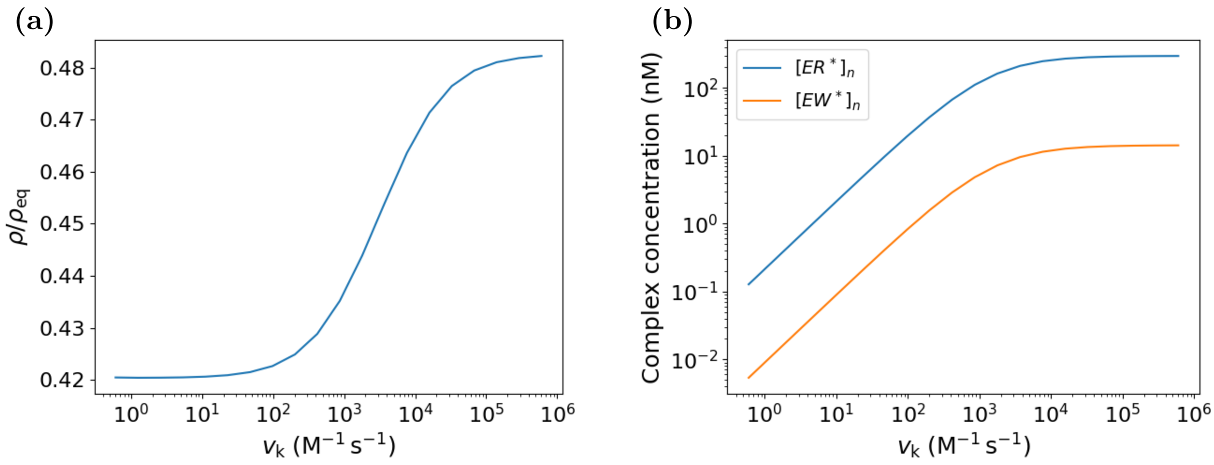

**Figure S6:** **(a)** Normalized specificity $\rho/\rho_{\mathrm{eq}}$ as a function of the kinase activation rate $v_k$. **(b)** Steady-state concentrations of the complexes $[ER^*]_n$ and $[EW^*]_n$ in the readout compartment as a function of $v_k$. The remaining parameters are fixed to the reference values reported in Sec. 2.

The slight increase of $\rho/\rho_{\mathrm{eq}}$ observed at large $v_k$ can be interpreted as a consequence of the enhanced activation flux. When kinase activity is very high, more activated wrong substrate $W^*$ is produced and transported through the domain. At fixed phosphatase rate $v_p$, this can make deactivation relatively less effective and slightly increase the contribution of $EW^*$ in the readout compartment.

In summary, these results indicate that $v_p$ and $v_k$ play distinct roles. The phosphatase rate $v_p$ controls the spatial confinement and lifetime of activated substrates during transport and therefore has a direct impact on proofreading. Instead, the kinase rate $v_k$ mainly controls the strength of the activation source and has only a secondary effect on discrimination in the parameter regime considered here.

##### 4.3. Effect of system size and diffusive transport time

We next analyzed the effect of the system size $L$, which sets the distance between the activation region and the readout region. Since the characteristic diffusive transport time scales as $\tau_D \sim L^2/D$, increasing $L$ is expected to enlarge the temporal window during which enzyme–substrate complexes can dissociate before reaching the readout compartment. We first performed a sweep over a range compatible with cellular length scales, $L \in [5, 21]\,\mu$m [3].

As shown in Fig. S7a, $\rho/\rho_{\mathrm{eq}}$ decreases as $L$ increases. This behavior is consistent with the spatial proofreading mechanism: a larger distance between the activation region and the readout region increases the effective transport time, thereby enhancing the probability that the wrong complex $EW^*$ dissociates before reaching the final compartment. As a result, the wrong substrate is penalized more strongly and discrimination improves. This interpretation is supported by the behavior of the complex concentrations shown in Fig. S7b. Both $[ER^*]_n$ and $[EW^*]_n$ decrease as $L$ increases, reflecting the weaker coupling between the activation source and the readout compartment, but the wrong complex decreases more steeply than the correct one. Consequently, the improvement in $\rho/\rho_{\mathrm{eq}}$ arises from a selective depletion of $EW^*$ relative to $ER^*$ at the readout.

Within the biologically plausible range explored in Fig. S7a, the dependence on $L$ is monotonic and does not display the non-monotonic minimum observed in Fig. 3a of the main text, where $\rho/\rho_{\mathrm{eq}}$ was plotted as a function of $\tau_D$ by varying $D$ at fixed $L$. However, this does not imply that $L$ and $D$ control proofreading through distinct mechanisms. In fact, the values of $L$ explored in this initial sweep correspond to a range of diffusive times that remains below the regime where the minimum becomes apparent. To test whether the relevant control parameter is the diffusive

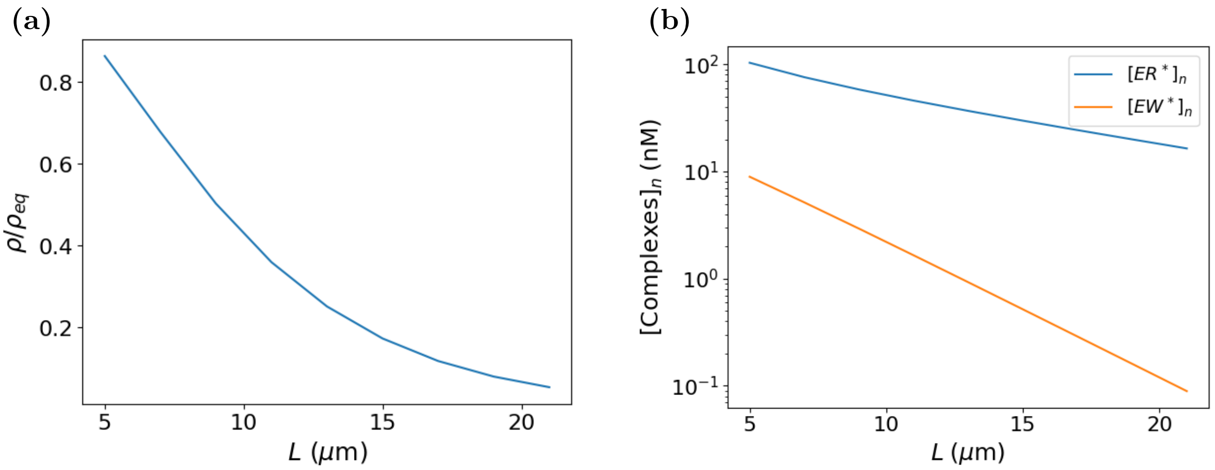

**Figure S7:** **(a)** Normalized specificity $\rho/\rho_{\mathrm{eq}}$ as a function of the system size $L$. **(b)** Steady-state concentrations of the complexes $[ER^*]_n$ and $[EW^*]_n$ in the readout compartment as a function of the system size $L$. The remaining parameters are fixed to the reference values reported in Sec. 2.

transport time, we compared simulations in which $\tau_D$ was varied either by changing $D$ at fixed $L$ or by changing $L$ at fixed $D$ (Fig. S8). The two sets of results approximately collapse over a broad range and display a minimum at comparable values of the transport time. This shows that the dominant control parameter is the diffusive transport time, which sets the temporal window available for selective dissociation before readout. Small deviations become visible only at the largest values of $\tau_D$, where the readout compartment is weakly supplied by activated species and the complex concentrations become very small. This indicates that the absence of a minimum in the initial $L$ sweep is due to the limited range of diffusive times accessible for cellular-scale values of $L$, rather than to a qualitative difference between varying $L$ and varying $D$.

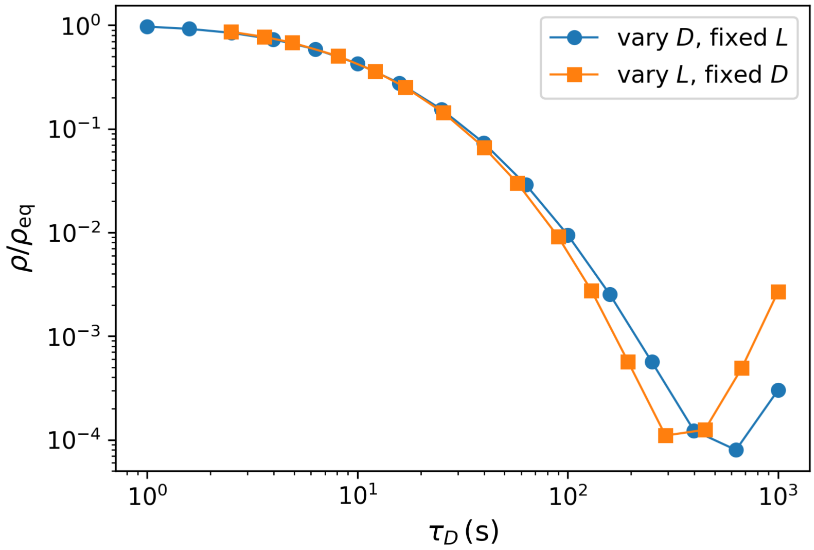

**Figure S8:** Normalized specificity $\rho/\rho_{\mathrm{eq}}$ as a function of the diffusive transport time $\tau_D = L^2/D$. Blue circles: simulations in which $\tau_D$ is varied by changing the diffusion coefficient $D$ at fixed system size $L = 10\,\mu\mathrm{m}$, with $D \in [10^{-1}, 10^{2}]\,\mu\mathrm{m}^2\,\mathrm{s}^{-1}$. Orange squares: simulations in which $\tau_D$ is varied by changing the system size $L$ at fixed diffusion coefficient $D = 10\,\mu\mathrm{m}^2\,\mathrm{s}^{-1}$, with $L$ chosen so as to span a comparable range of diffusive times. The remaining parameters are fixed to the reference values reported in Sec. 2.

##### 4.4. Effect of reduced complex diffusivity

In the reference model, all species are assumed to diffuse with the same coefficient $D$. From a physical perspective, this is only an approximation, since complex formation increases the hydrodynamic size of the diffusing species and is therefore expected to reduce its diffusivity. To test the robustness of the results with respect to this simplification, we considered a variant of the model in which the free species have diffusion coefficient $D$, while the complexes diffuse more slowly:

$$D_{\mathrm{c}} = \gamma D, \qquad \gamma = \tfrac{1}{2},$$

where $D_{\mathrm{c}}$ denotes the common diffusion coefficient of the complexes $ER^*$ and $EW^*$. This modification introduces a distinct diffusive transport time for the complexes, $\tau_D^{\mathrm{c}} = L^2/\gamma D$, which is larger than the transport time of the free species.

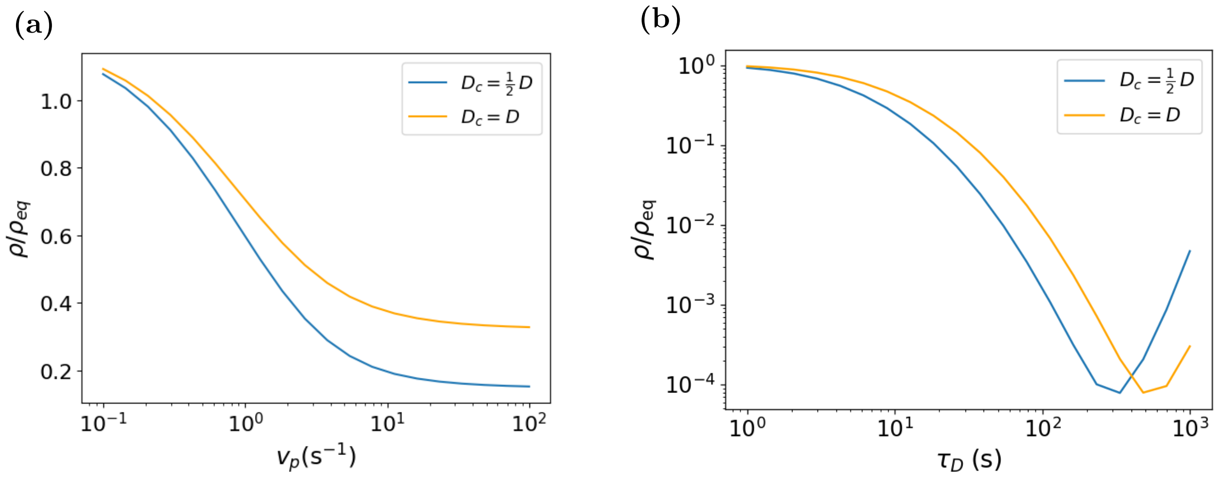

**Figure S9:** Comparison between the reference model with uniform diffusion, $D_c = D$, and a model with reduced complex diffusivity, $D_c = \frac{1}{2}D$. **(a)** Logarithmic sweep of the phosphatase rate $v_p$. **(b)** Normalized specificity as a function of the nominal diffusive transport time $\tau_D$; the system size is kept fixed at $L = 10\,\mu$m, while the nominal diffusion coefficient is varied in the range $D \in [0.1, 100]\,\mu\mathrm{m}^2\,\mathrm{s}^{-1}$. The remaining parameters are fixed to the reference values reported in Sec. 2.

Figure S9a shows that reducing the diffusion coefficient of the complexes leads to smaller values of $\rho/\rho_{\mathrm{eq}}$ at fixed $v_p$, corresponding to improved discrimination. This behavior is consistent with the spatial proofreading mechanism: slower complex diffusion increases the complex transport time $\tau_D^{\mathrm{c}}$, thereby enlarging the time available for selective dissociation during transport. As a result, wrong complexes are more strongly depleted before reaching the readout compartment, whereas correct complexes are comparatively less affected because $k_{\mathrm{off}}^W > k_{\mathrm{off}}^R$.

Moreover, reducing the diffusivity of the activated complexes preserves the qualitative non-monotonic dependence of the normalized specificity on $\tau_D$, while shifting the minimum toward smaller values (Fig. S9b). This shift is expected because, at fixed $\tau_D$, the effective transport time of the complexes is longer when $D_c = D/2$. It therefore reflects the need for a larger nominal diffusion coefficient $D$ to offset the slower transport of the activated complexes. This behavior is consistent with the interpretation of the minimum as a crossover between a transport-limited regime and a regime controlled by local reaction kinetics.

Overall, these results show that the assumption $D_c = D$ does not alter the qualitative physics of the model, although it affects the quantitative level of discrimination and the location of the optimal transport regime. The enhancement of discrimination associated with slower transport does not require a global reduction of the diffusivity of all species; it is sufficient that the complexes diffuse more slowly, since they directly determine the readout signal. For simplicity, the reference model adopts the choice $D_c = D$, which reduces the number of parameters while retaining the essential mechanism of spatial proofreading.

#### 5. Additional two-dimensional simulations

##### 5.1. Kinase localization patterns in rectangular domains

To further test the role of kinase spatial organization, we compared different kinase configurations within the same rectangular domain, while keeping the total catalytic area fixed. In the first configuration, kinase activity was distributed in several active bands along the source boundary, whereas in the second configuration the same total active length was concentrated into a single central band. Representative examples of the two configurations are shown in Fig. S10. The specificity was measured along the opposite boundary, which defines the readout boundary.

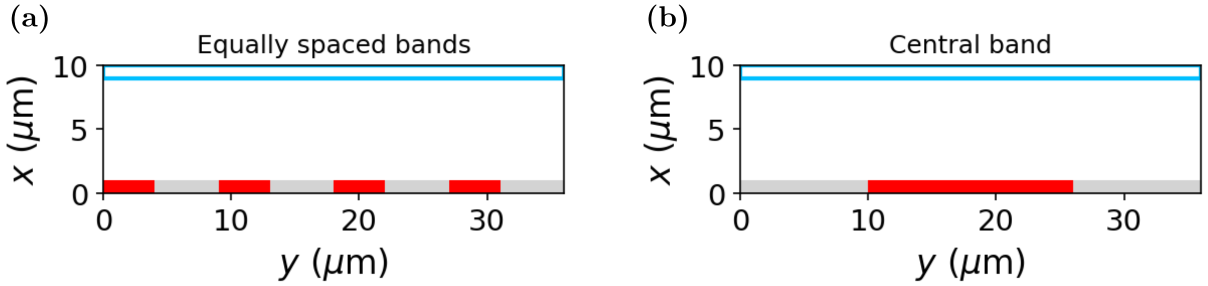

**Figure S10:** Examples of rectangular domains of size $W = 10\,\mu$m and $H = 36\,\mu$m, discretized with $\Delta x = \Delta y = 1\,\mu$m. Along the source boundary, red and grey cells indicate compartments with and without kinase activity, respectively. The cyan line marks the readout boundary at the opposite side of the domain. **(a)** Distributed kinase pattern with active bands of width $L_y = 4\,\mu$m separated by $d = 5\,\mu$m. **(b)** Single central kinase band of width $L_{\text{CB}} = 16\,\mu$m.

The two geometries lead to qualitatively different profiles of the normalized specificity along the readout boundary (Fig. S11). In the distributed-band geometry, $\rho(y)/\rho_{\text{eq}}$ remains close to the reference value, with weak oscillations that become more pronounced as the distance $d$ between neighboring kinase bands increases. This occurs because larger separations reduce the lateral overlap between the concentration gradients generated by different bands. In contrast, the central-band geometry produces a more heterogeneous profile: the normalized specificity is largest in the region aligned with the kinase band and decreases toward the lateral edges of the readout boundary. These edge regions are associated with longer effective transport paths from the activation source, providing a larger temporal window for selective dissociation and phosphatase-mediated deactivation.

The heatmaps in Fig. S12 provide a spatial visualization of this mechanism. In the central-band geometry, both activated substrates, $R^*$ and $W^*$, decrease with distance from the kinase band, but the depletion is stronger for the wrong activated substrate $W^*$. The same effect is even more evident at the level of the enzyme–substrate complexes: $EW^*$ decays more rapidly than $ER^*$ along the diffusive paths connecting the source to the readout boundary. This reflects the lower stability of the wrong branch, since wrong complexes dissociate more frequently during transport; once dissociated, $W^*$ is exposed again to phosphatase-mediated deactivation and is therefore less likely to persist up to the readout region. Instead, distributed bands generate overlapping gradients that smooth these spatial variations, leading to a weaker modulation of the normalized specificity.

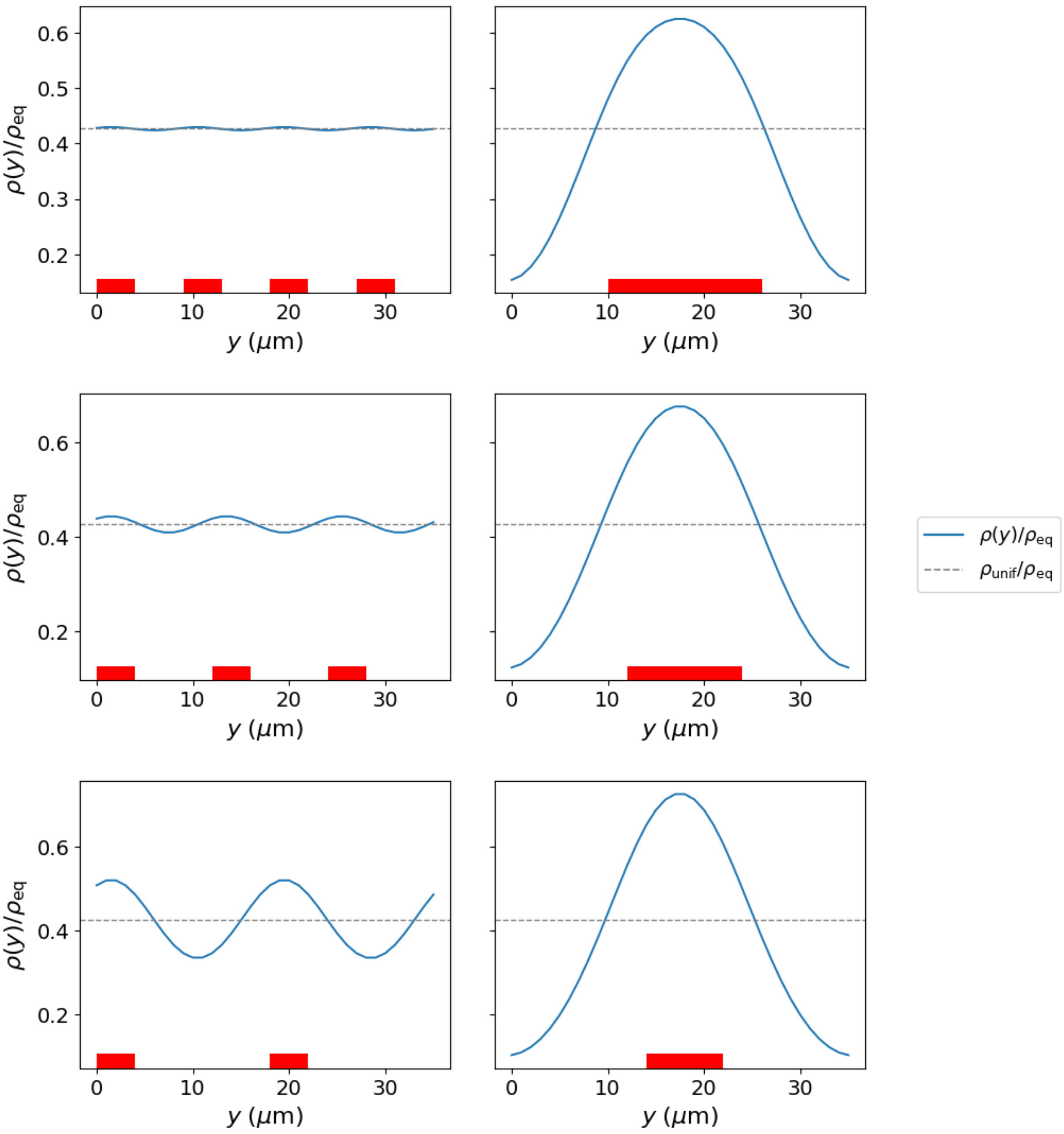

**Figure S11:** Normalized specificity profiles $\rho(y)/\rho_{\mathrm{eq}}$ along the readout boundary for distributed kinase bands (left column) and for a single central kinase band (right column). The three rows correspond to distributed-band separations $d = 5, 8, 14\,\mu$m, respectively. The red regions shown at the bottom of each panel indicate the kinase-active portions of the source boundary. The local kinase rate $v_k$ is adjusted in each configuration so that the total stationary activation flux is the same across all source geometries. Dashed lines indicate the normalized reference value obtained from a uniform kinase distribution normalized according to the same criterion, corresponding to $\rho_{\mathrm{unif}}/\rho_{\mathrm{eq}} \simeq 0.43$. The rectangular domain has size $W = 10\,\mu$m and $H = 36\,\mu$m, with spatial discretization $\Delta x = \Delta y = 1\,\mu$m. Distributed bands have width $L_y = 4\,\mu$m. The corresponding central-band lengths are $L_{\mathrm{CB}} = 16, 12, 8\,\mu$m, respectively, chosen so as to conserve the total catalytic area. The remaining parameters are those used in Fig. 2 of the main text.

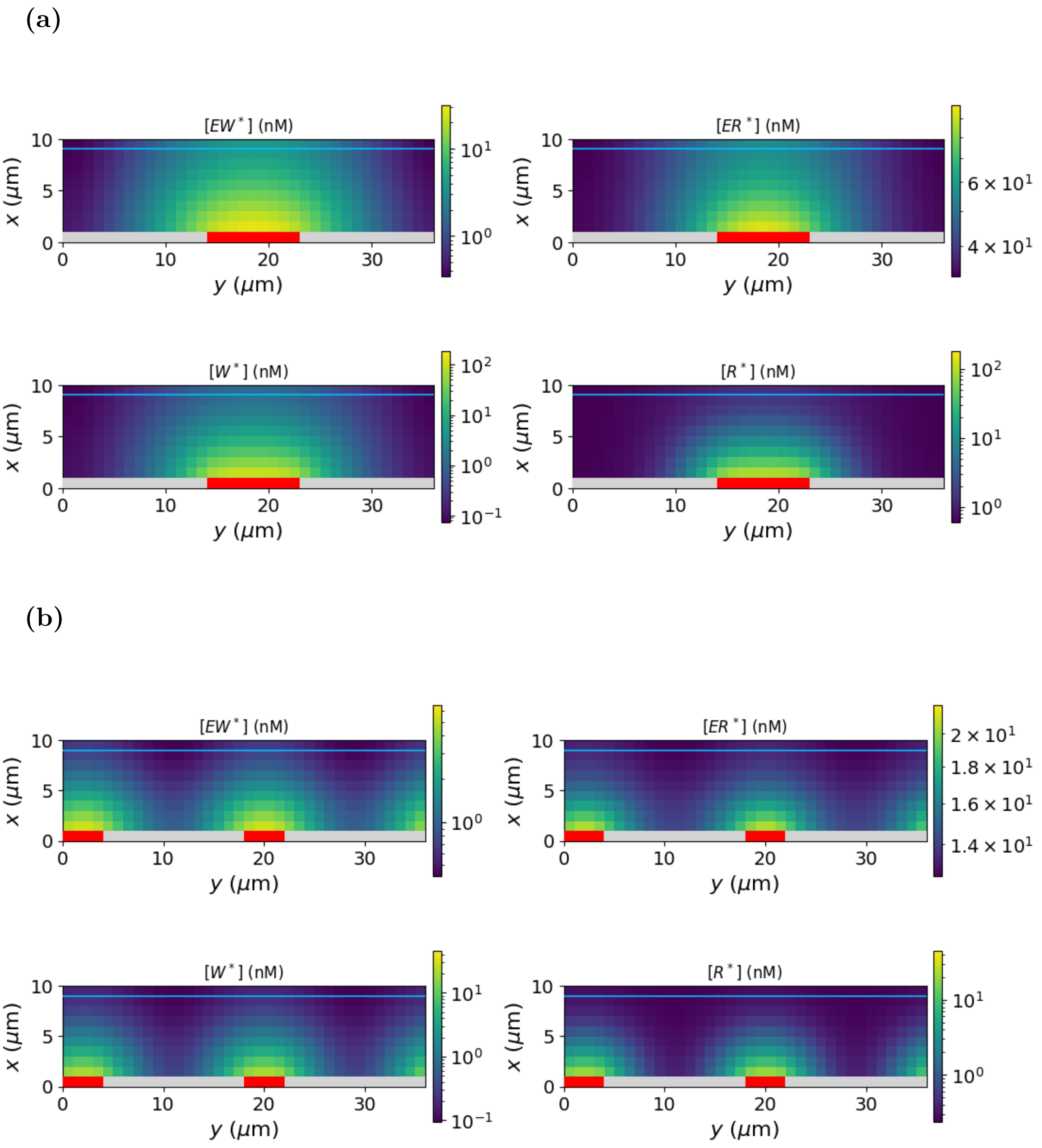

**Figure S12:** Spatial distributions of activated substrates and complexes in the rectangular domain. **(a)** Single central kinase band with $L_{\mathrm{CB}} = 9\,\mu$m. **(b)** Distributed-band geometry with band width $L_y = 4\,\mu$m and separation $d = 14\,\mu$m. In both cases, the domain has size $W = 10\,\mu$m and $H = 36\,\mu$m, with $\Delta x = \Delta y = 1\,\mu$m. In the first compartment, red/grey cells indicate the presence/absence of kinases. The cyan line separates the last compartment, identified as the readout region. The remaining parameters are fixed to the reference values reported in Sec. 2.

##### 5.2. Effect of the central-band length

The comparison between distributed and central kinase patterns shows that a single central activation band generates a strongly heterogeneous readout profile, whereas distributed bands lead to a much smoother modulation along the boundary. We therefore analyzed the central-band geometry in more detail by varying the band length $L_{\mathrm{CB}}$ and measuring the resulting specificity profile $\rho(y)/\rho_{\mathrm{eq}}$. The resulting profiles are shown in Fig. S13.

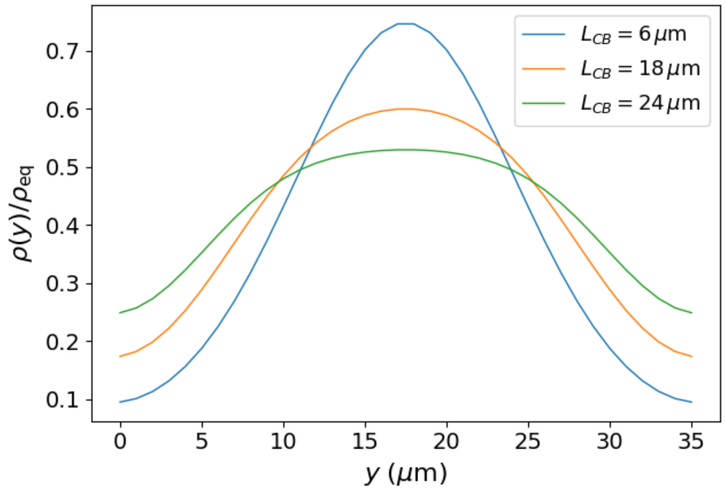

**Figure S13:** Specificity profiles $\rho(y)/\rho_{\mathrm{eq}}$ along the readout boundary for different values of the central-band length $L_{\mathrm{CB}}$. The rectangular domain has size $W = 10\,\mu$m and $H = 36\,\mu$m, with $\Delta x = \Delta y = 1\,\mu$m. The remaining parameters are fixed to the reference values reported in Sec. 2.

As $L_{\mathrm{CB}}$ increases, the profile becomes progressively more homogeneous. The central peak in $\rho(y)/\rho_{\mathrm{eq}}$ is reduced and broadened, while the low-specificity regions near the lateral edges become less pronounced. This trend is consistent with the geometric interpretation of the two-dimensional results. A short central band creates a broad range of effective distances from the activation source: readout points close to the center are directly aligned with the kinase region, whereas points near the lateral edges are reached only after a longer transverse diffusive path. This enhances the spatial proofreading effect at the edges, where the wrong complex is more strongly depleted before readout. Increasing $L_{\mathrm{CB}}$ reduces this transverse distance heterogeneity and drives the system toward the uniformly activated limit.

To quantify this effect, we focused on the corners of the readout boundary, where the minimum values of $\rho(y)/\rho_{\mathrm{eq}}$ are observed for narrow central bands. We define the normalized corner specificity

$$\frac{\rho_{\mathrm{c}}(L_{\mathrm{CB}})}{\rho_{\mathrm{c}}(H)}, \tag{S21}$$

where $\rho_{\mathrm{c}}(L_{\mathrm{CB}})$ is the specificity measured at the corner of the readout boundary for a central band of length $L_{\mathrm{CB}}$ and $\rho_{\mathrm{c}}(H)$ is the corresponding value in the uniformly activated limit, $L_{\mathrm{CB}} = H$. The result is shown in Fig. S14.

The normalized corner specificity decreases markedly as the central band becomes shorter, reaching values around 0.2 for narrow bands. Thus, localizing the kinase in a sufficiently small central region can substantially enhance discrimination at readout positions that are far from the activation source. Conversely, as $L_{\mathrm{CB}}/H$ approaches one, the corner specificity approaches the uniformly activated case. This confirms that the spatial organization of the kinase does not only affect the average proofreading performance, but also redistributes discrimination along the readout boundary.

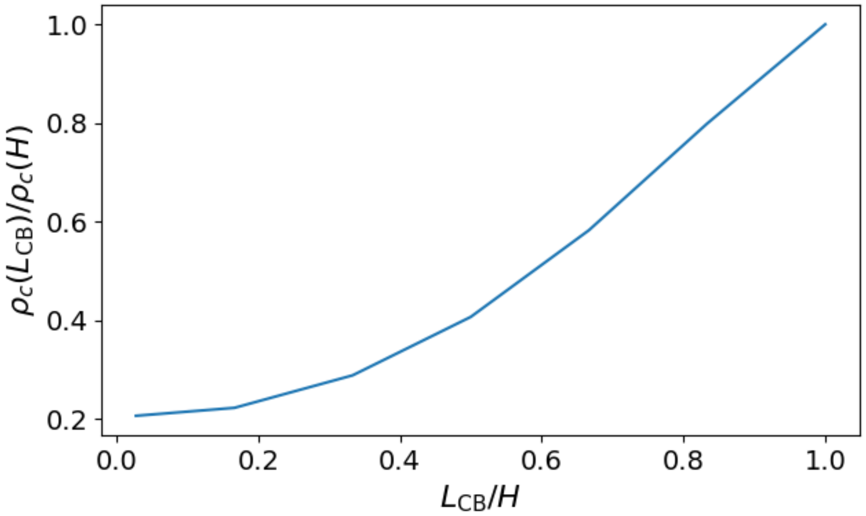

**Figure S14:** Normalized corner specificity $\rho_c(L_{CB})/\rho_c(H)$ as a function of $L_{CB}/H$. The rectangular domain has size $W = 10\,\mu$m and $H = 36\,\mu$m, with $\Delta x = \Delta y = 1\,\mu$m. The remaining parameters are fixed to the reference values reported in Sec. 2.

##### 5.3. Effect of the diffusion coefficient $D$

Finally, we analyzed how diffusion affects the readout profile in the two-dimensional central-band geometry. Since the domain width $W$ is kept fixed, varying the diffusion coefficient $D$ is equivalent to varying the nominal diffusive transport time $\tau_D = W^2/D$ between the activation boundary and the readout boundary. This provides a direct connection with the one-dimensional results, while retaining the additional spatial heterogeneity introduced by the central-band geometry. We focus on this geometry because it generates a strongly heterogeneous readout profile and therefore makes the interplay between transport and spatial organization more visible. The resulting normalized specificity profiles along the readout boundary are shown in Fig. S15.

For large values of $D$, diffusion rapidly mixes the activated species across the domain and the normalized specificity profile becomes relatively homogeneous, approaching the weak-discrimination regime. As $D$ is decreased, the effective transport time increases and the spatial proofreading mechanism becomes more efficient: $\rho(y)/\rho_{eq}$ decreases over most of the readout boundary, indicating improved discrimination. In this regime, the profile retains the same qualitative structure observed at the reference value of $D$, with a maximum close to the region aligned with the central activation band and lower values toward the lateral edges.

At the smallest value considered, $D = 0.1\,\mu\text{m}^2\,\text{s}^{-1}$, the normalized specificity profile changes qualitatively. It is no longer minimized at the edges; instead, $\rho(y)/\rho_{eq}$ increases away from the central region. This behavior is consistent with the one-dimensional analysis at very long transport times. When diffusion is too slow, the readout boundary becomes weakly supplied with activated species from the kinase region. In this regime, local complex formation in the readout compartment becomes relatively more important than the diffusive influx from the activation source and the spatial proofreading advantage is consequently reduced.

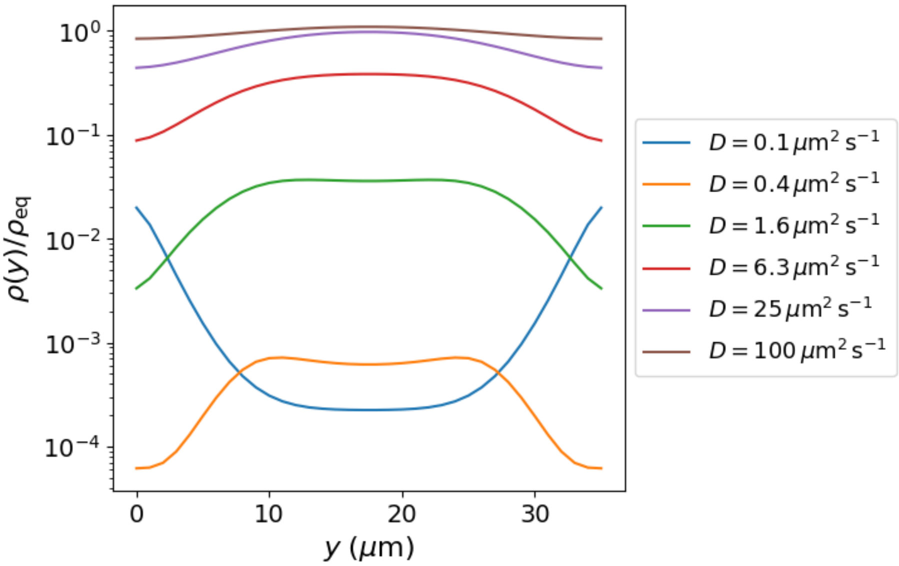

**Figure S15:** Normalized specificity profiles $\rho(y)/\rho_{\mathrm{eq}}$ along the readout boundary for different values of the diffusion coefficient $D$ in the central-band geometry. The rectangular domain has size $W = 10\,\mu\mathrm{m}$ and $H = 36\,\mu\mathrm{m}$, with $\Delta x = \Delta y = 1\,\mu\mathrm{m}$. The central kinase band has length $L_{\mathrm{CB}} = 18\,\mu\mathrm{m}$. The remaining parameters are fixed to the reference values reported in Sec. 2.

To clarify this change of regime, we compared the diffusive influx contribution and the local formation term of $EW^*$ along the readout boundary for $D = 10\,\mu\mathrm{m}^2\,\mathrm{s}^{-1}$ and $D = 0.1\,\mu\mathrm{m}^2\,\mathrm{s}^{-1}$, as shown in Fig. S16. For $D = 10\,\mu\mathrm{m}^2\,\mathrm{s}^{-1}$, the diffusive contribution remains larger than the local formation term over the readout boundary, indicating that the wrong-complex concentration is mainly controlled by transport from the activation region. By contrast, for $D = 0.1\,\mu\mathrm{m}^2\,\mathrm{s}^{-1}$, the local formation term dominates over the diffusive contribution. In this limit, the spatially transported flux is too weak to maintain the same selective depletion mechanism and the normalized specificity profile is instead controlled primarily by local readout kinetics.

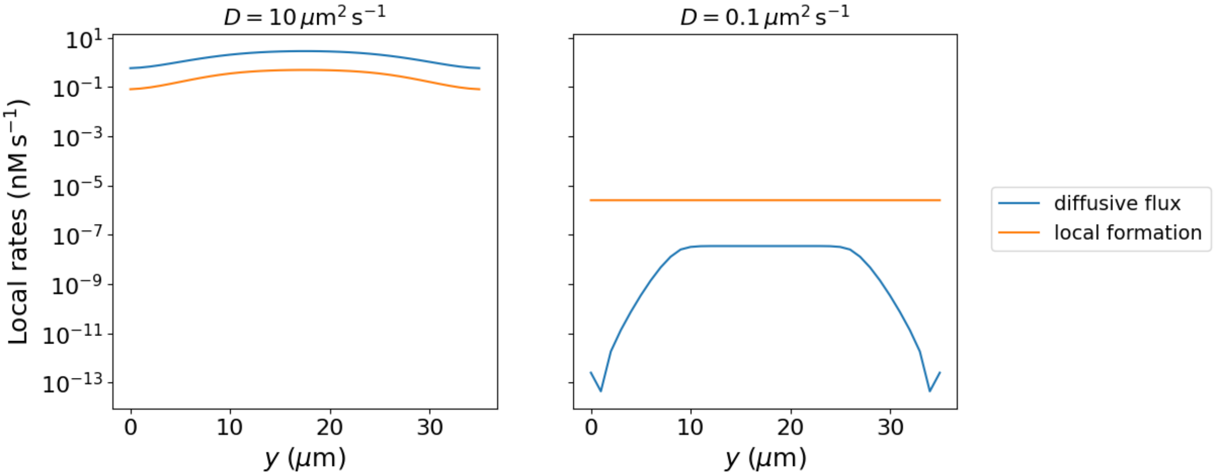

**Figure S16:** Comparison between the diffusive influx contribution and the local formation term of $EW^*$ along the readout boundary in the central-band geometry for $D = 10\,\mu\mathrm{m}^2\,\mathrm{s}^{-1}$ and $D = 0.1\,\mu\mathrm{m}^2\,\mathrm{s}^{-1}$. The rectangular domain has size $W = 10\,\mu\mathrm{m}$ and $H = 36\,\mu\mathrm{m}$, with $\Delta x = \Delta y = 1\,\mu\mathrm{m}$, and the central kinase band has length $L_{\mathrm{CB}} = 18\,\mu\mathrm{m}$. The remaining parameters are fixed to the reference values reported in Sec. 2.

Overall, this two-dimensional diffusion sweep confirms the interpretation obtained in one dimension. Decreasing $D$ initially improves discrimination because it increases the transport time available for selective dissociation. However, when diffusion becomes too slow, the readout region is no longer efficiently supplied by activated species from the source and local formation terms become dominant. Thus, also in two dimensions, proofreading is optimized at intermediate transport conditions: diffusion must be slow enough to allow selective dissociation during transport, but not so slow that the readout becomes dominated by local formation kinetics.

#### 6. Activity–specificity trade-off

An important issue in proofreading systems is the possible trade-off between discrimination and readout activity. In our model, proofreading performance is quantified through the normalized specificity $\rho/\rho_{\mathrm{eq}}$, while activity is quantified by the steady-state concentration of the correct complex in the readout compartment,

$$A \equiv [ER^*]_n. \tag{S22}$$

Experimentally, activity is an important constraint in the implementation of proofreading circuits, since the concentration of the correct complex must be large enough to be detectable and to elicit a downstream response. Biologically, a sufficiently large activity is also required to distinguish the proofreading output from molecular fluctuations and background noise [5]. Therefore, it is important to verify whether the parameter regimes in which the normalized specificity reaches low values are also associated with a sufficiently large concentration of the correct complex. Following Ref. [5], we use $A = 1\,\mathrm{nM}$ as a reference activity benchmark. This value should not be interpreted as a universal biological limit, but rather as a useful benchmark to identify regimes in which strong discrimination is accompanied by a non-negligible correct readout.

As a first control, we investigated how proofreading performance depends on the initial substrate concentration. This is particularly relevant in view of the results reported in Ref. [5]. In their intracellular mass-conserving spatial proofreading model, increasing the initial substrate concentrations up to $10\,\mu\mathrm{M}$ restores a sufficiently large activity but strongly reduces discrimination. We therefore varied the initial substrate concentration $[S_0]$, keeping $[R_0] = [W_0]$, under two different conditions: first, at fixed enzyme concentration $[E_0]$, and second, at fixed enzyme-to-substrate ratio $[E_0]/[S_0]$.

Figure S17a shows that the dependence of the normalized specificity on the initial substrate concentration is generally weak over a broad range of $[S_0]$. In particular, when $[E_0]$ is kept fixed, $\rho/\rho_{\mathrm{eq}}$ remains approximately constant. By contrast, when the ratio $[E_0]/[S_0]$ is kept fixed, the normalized specificity progressively increases at large $[S_0]$. This behavior is consistent with the effect observed in the sweep of $[E_0]$ discussed in Sec. 4.1, since increasing $[S_0]$ at fixed $[E_0]/[S_0]$ also increases the total enzyme concentration and therefore favors rebinding.

The corresponding activity is shown in Fig. S17b. At the reference concentration $[S_0] = 3\,\mu\mathrm{M}$, the correct-complex concentration at the readout is well above the $1\,\mathrm{nM}$ reference activity benchmark, while the normalized specificity remains substantially below unity. Therefore, the reference choice of $[S_0]$ does not correspond to a regime in which enhanced discrimination is obtained at the cost of a vanishingly small correct readout.

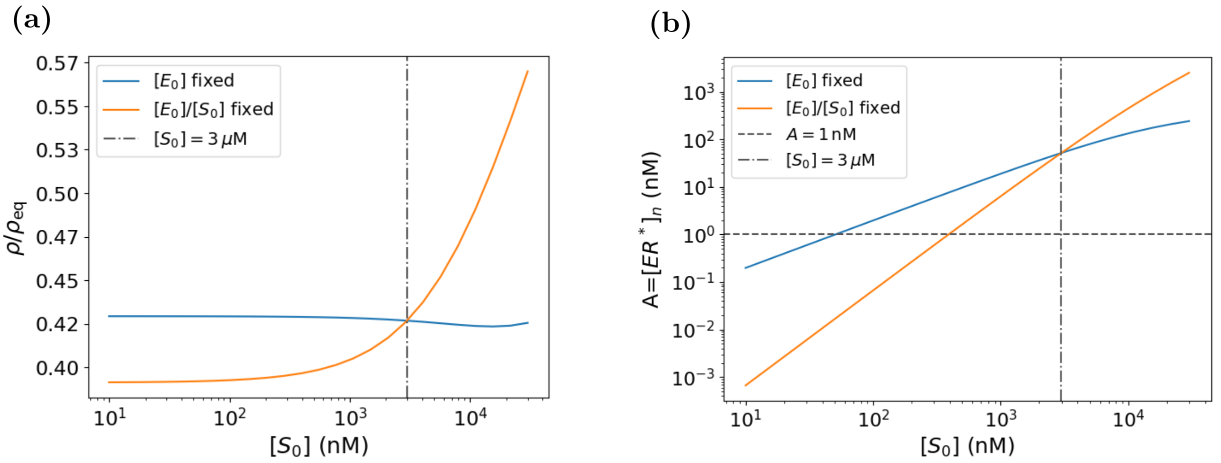

**Figure S17:** **(a)** Normalized specificity $\rho/\rho_{\text{eq}}$ as a function of $[S_0]$ (with $[R_0] = [W_0]$), either keeping $[E_0] = 0.6\,\mu\text{M}$ fixed or keeping $[E_0]/[S_0] = 0.2$ fixed. **(b)** Corresponding activity $A = [ER^*]_n$ in the readout compartment. The horizontal dashed line in panel **(b)** indicates the reference activity benchmark $A = 1\,\text{nM}$, while the vertical dash-dotted line in both panels marks the reference concentration $[S_0] = 3\,\mu\text{M}$. The remaining parameters are fixed to the reference values reported in Sec. 2.

We next asked whether the parameter regimes explored in the main text that strongly affect proofreading performance also retain a sufficiently large readout activity. We focus on the phosphatase rate $v_p$, the diffusive transport time $\tau_D$, and the absolute scale of the dissociation rates. For each sweep, we examined the steady-state concentrations of the correct and wrong complexes in the readout compartment and compared the correct-complex activity with the reference benchmark. The results are shown in Fig. S18.

For $v_p$, the correct-complex activity remains above the 1 nM benchmark over the entire range explored. Therefore, the improvement in normalized specificity observed in Fig. 2b is not associated with a depletion of the correct readout.

A different behavior is observed when varying $\tau_D$: at sufficiently large values, corresponding to slow diffusion, the activity progressively decreases and eventually falls below the reference benchmark. The minimum in normalized specificity is reached close to this low-activity region (see the orange curve in Fig. 3a), indicating that an activity–specificity compromise arises when transport becomes sufficiently slow. However, substantial proofreading gains are already obtained at shorter transport times, where the correct-complex concentration remains well above the benchmark. Moreover, at still larger $\tau_D$ the activity continues to decrease while the normalized specificity increases again toward equilibrium. Thus, reducing the readout activity indefinitely does not produce a corresponding indefinite improvement in discrimination.

A similar distinction emerges from the sweep of the dissociation rates at fixed $\alpha = k_{\text{off}}^W/k_{\text{off}}^R$. The activity decreases strongly for $k_{\text{off}}^R \gtrsim 1\,\text{s}^{-1}$ and eventually falls below the reference benchmark. However, comparison with the corresponding $\alpha = 10$ specificity curve (red curve in Fig. 5b) shows that the minimum of $\rho/\rho_{\text{eq}}$ is reached before entering this strongly activity-depleted regime. Further increasing $k_{\text{off}}^R$ reduces the activity while simultaneously worsening discrimination, with $\rho/\rho_{\text{eq}}$ approaching unity. Hence, in this case the minimum normalized specificity cannot be attributed simply to a vanishing correct-complex concentration.

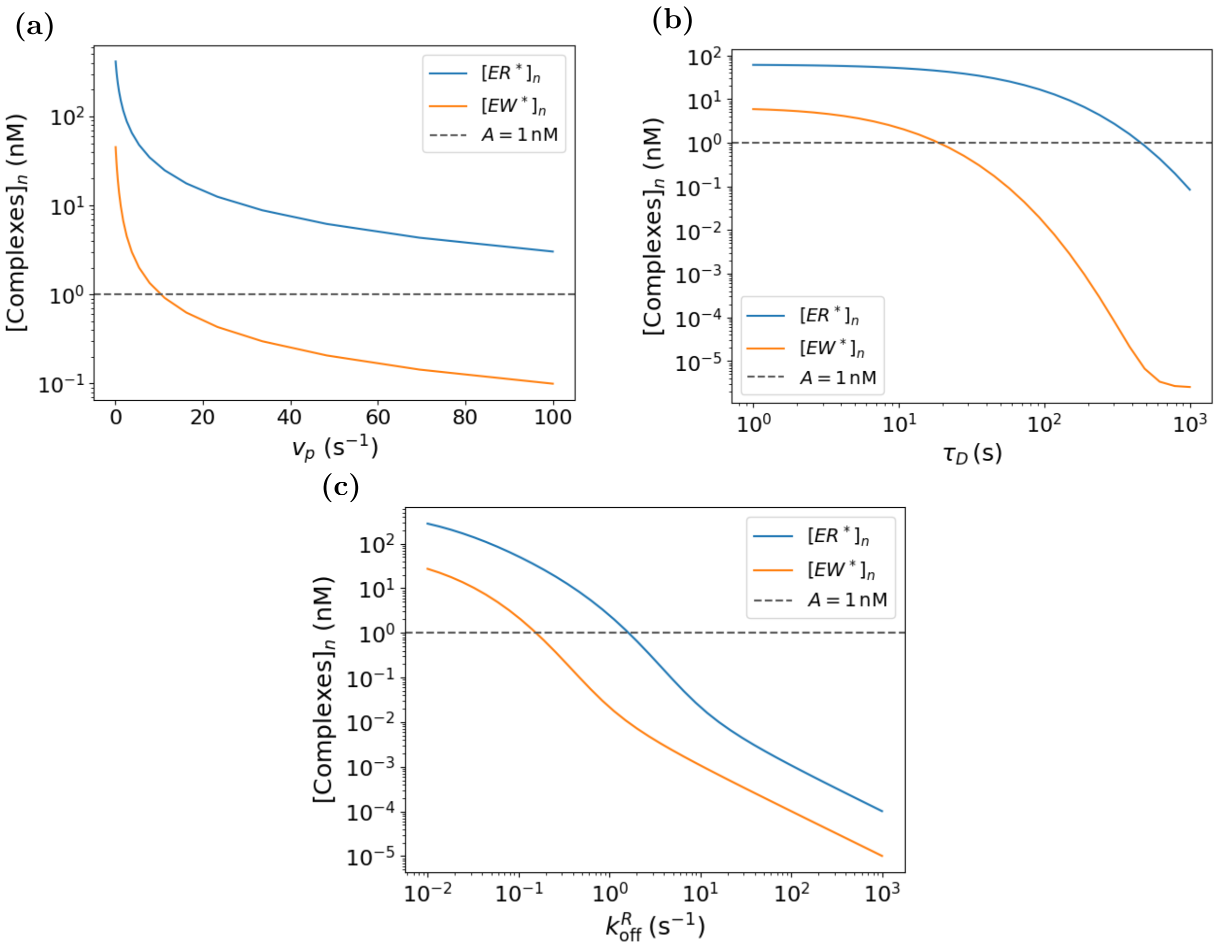

**Figure S18:** Steady-state concentrations of the correct and wrong complexes, $[ER^*]_n$ and $[EW^*]_n$, in the readout compartment for the main kinetic and transport parameter sweeps. **(a)** Dependence on the phosphatase rate $v_p$. **(b)** Dependence on the diffusive transport time $\tau_D$, obtained by varying $D$ at fixed $L = 10\,\mu$m. **(c)** Dependence on the correct-complex dissociation rate $k^R_{\rm off}$, while keeping the affinity ratio $\alpha = k^W_{\rm off}/k^R_{\rm off} = 10$ fixed. The horizontal dashed line indicates the reference activity benchmark $A = 1\,$nM. The remaining parameters are fixed to the reference values reported in Sec. 2.

As a final control, we performed a random sampling of the parameter space defined by the ranges of $D$, $v_p$, and $k^R_{\rm off}$ considered in the previous sweeps. For each randomly sampled parameter set, we computed the steady-state correct-complex activity $A = [ER^*]_n$ and the corresponding normalized specificity $\rho/\rho_{\rm eq}$, and represented the results in the activity–specificity plane. This analysis allows us to determine whether strong proofreading is systematically restricted to regimes of very low activity, or whether substantial discrimination can coexist with a correct-complex concentration above the reference activity benchmark.

The resulting activity–specificity landscape is shown in Fig. S19. The sampled parameter sets span a broad region of the plane, including both low- and high-activity regimes. Notably, many configurations lie in the region $A > 1\,$nM and $\rho/\rho_{\rm eq} < 1$, showing that enhanced discrimination is not restricted to parameter sets characterized by a strongly depleted correct readout. At the same time, some parameter combinations lead to activities below the reference benchmark, consistently with the behavior observed in the individual sweeps of Fig. S18. Thus, within the explored parameter ranges, an activity–specificity compromise can arise in specific regions of parameter space, but strong proofreading gains can also be achieved while maintaining an appreciable correct-complex concentration.

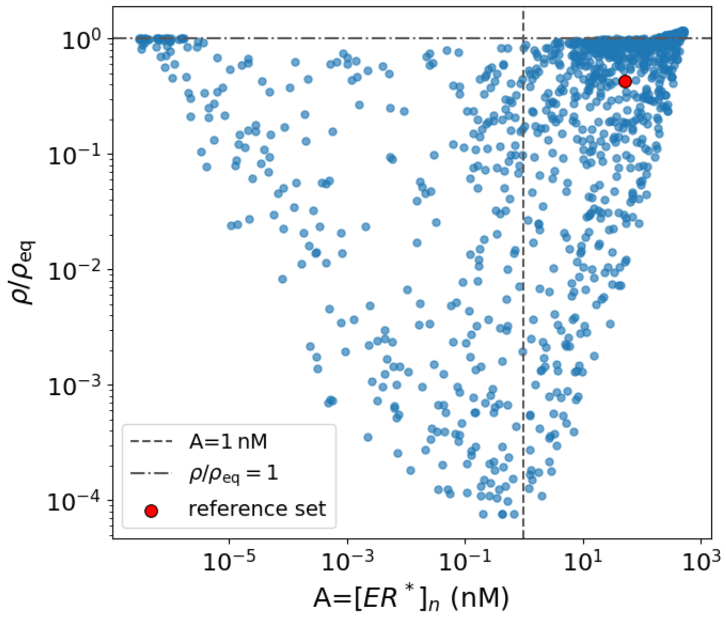

**Figure S19:** Global activity–specificity landscape obtained from 1000 independently sampled parameter sets, with $D$, $v_p$, and $k^R_{\text{off}}$ sampled uniformly on a logarithmic scale over the ranges $D \in [10^{-1}, 10^2]\,\mu\text{m}^2\,\text{s}^{-1}$, $v_p \in [10^{-1}, 10^2]\,\text{s}^{-1}$, and $k^R_{\text{off}} \in [10^{-2}, 10]\,\text{s}^{-1}$, while keeping the affinity ratio fixed at $\alpha = 10$. The vertical dashed line indicates the reference activity benchmark $A = 1\,\text{nM}$, while the horizontal dash-dotted line marks the equilibrium value $\rho/\rho_{\text{eq}} = 1$. The remaining parameters, as well as the reference parameter set highlighted by a red marker, are defined in Sec. 2.

Overall, these analyses show that enhanced proofreading is not intrinsically associated with a vanishingly small correct readout. Although an activity–specificity compromise emerges in some regions of parameter space, substantial discrimination can be achieved while maintaining an appreciable activity above the reference benchmark.
